# Limited time window for retinal gene therapy in a preclinical model of ciliopathy

**DOI:** 10.1101/2020.05.05.072793

**Authors:** Poppy Datta, Avri Ruffcorn, Seongjin Seo

## Abstract

Retinal degeneration is a common clinical feature of ciliopathies, a group of genetic diseases linked to ciliary dysfunction, and gene therapy is an attractive treatment option to prevent vision loss. Although the efficacy of retinal gene therapy is well established by multiple proof-of-concept preclinical studies, its long-term effect, particularly when treatments are given at advanced disease stages, is controversial. Incomplete treatment and intrinsic variability of gene delivery methods may contribute to the variable outcomes. Here, we used a genetic rescue approach to “optimally” treat retinal degeneration at various disease stages and examined the long-term efficacy of gene therapy in a mouse model of ciliopathy. We used a Bardet-Biedl syndrome type 17 (BBS17) mouse model, in which the gene-trap that suppresses *Bbs17* (also known as *Lztfl1*) expression can be removed by tamoxifen administration, restoring normal gene expression systemically. Our data indicate that therapeutic effects of retinal gene therapy decrease gradually as treatments are given at later stages. These results suggest the presence of limited time window for successful gene therapy in certain retinal degenerations. Our study also implies that the long-term efficacy of retinal gene therapy may depend on not only the timing of treatment but also other factors such as the function of mutated genes and residual activities of mutant alleles.

## INTRODUCTION

Approximately a quarter of inherited retinal degeneration (IRD) genes are involved in building and maintaining the sensory cilium of photoreceptor cells, called the outer segment (OS) (1-4) (also see the RetNet database: https://sph.uth.edu/retnet/). Retinal degenerations caused by cilium-related defects, or retinal ciliopathies, are particularly detrimental because degeneration often begins very early in life (e.g. in infancy or early childhood) and ultimately results in severe vision loss. Gene therapy is a promising treatment option for IRDs including retinal ciliopathies. Preclinical studies conducted in various animal models of IRDs have shown that gene therapies improve functional vision and prevent or delay retinal degeneration (reviewed in (5, 6) and references therein). Furthermore, clinical trials of *RPE65* gene augmentation therapy conducted in Leber congenital amaurosis type 2 (LCA2) patients not only demonstrated the safety of adeno-associated virus (AAV)-mediated gene delivery but also led to a remarkable improvement in functional vision in some subjects (7-14). These positive outcomes excited the research community along with patient groups, and multiple clinical trials for various retinal disorders are currently underway (https://clinicaltrials.gov).

Although the efficacy of retinal gene therapies is well established when the treatment is given before the onset or at early stages of a disease, long-term effects of mid-to-late stage gene therapies are controversial (15-23). In most proof-of-concept preclinical studies, therapeutic genes were delivered before degeneration became evident. Although these studies demonstrate that gene therapies prevent retinal degeneration and preserve vision as long as therapeutic genes are delivered “early enough”, such experimental settings do not closely mimic the clinical settings of human diseases, as most patients are diagnosed after retinal degeneration has significantly progressed. Indeed, several follow-up studies of *RPE65* gene therapy clinical trials found that retinal degeneration continued despite initial and sustained vision improvement (15-17). Continued retinal degeneration was also observed in a canine model of *RPE65*-LCA, in which gene therapy vectors were delivered after the onset of retinal degeneration (15). In contrast, some other studies found that gene therapies after the onset of degeneration stopped further progression of degeneration (18-22). A recent study conducted in an *RPE65* dog model found that the long-term outcome of gene therapy was dependent on the health of the photoreceptor cell layer; gene therapy arrested retinal degeneration in areas where more than 63% of photoreceptors were remaining at the time of intervention, but retinal degeneration continued in areas with fewer retained photoreceptors (23).

Although the basis of these divergent outcomes is not well defined, technical limitations intrinsic to the current gene delivery methods are thought to be a contributing factor. Currently, subretinal injection of viral vectors such as AAV is the most broadly used gene delivery method. Subretinal injections are inherently variable and could themselves cause some injury to the retina. In addition, each subretinal injection treats only a part of the retina and the number of cells actually transduced varies from injection to injection. As a result, significant and variable proportions of cells in treated eyes usually remain untreated. Dying cells in the retina may cause secondary death of neighboring, successfully rescued cells. Another potential problem is insufficient, excessive, or unregulated expression of therapeutic genes. Since promoters and enhancers of most disease genes are not well defined and all viral vectors have a limited cargo capacity (e.g. ∼4.7 kb for AAV), it is difficult to use therapeutic genes’ natural promoters. Instead, small promoters such as cytomegalovirus (CMV) immediate-early promoter or photoreceptor-specific GRK1 promoter are usually used. Combined with the variability in the number of viral particles transduced into a cell, expression levels of therapeutic genes could be outside of optimal or even tolerable ranges.

In recent studies conducted in a mouse model of *PDE6B*-associated retinitis pigmentosa (RP), Tsang and colleagues circumvented the technical problems of subretinal injection by inducibly restoring the expression of endogenous *Pde6b* and compared the efficacy of gene therapies at various disease stages (24-26). These studies found that in “optimally treated” retinas (i.e. gene expression was restored to the endogenous level in the vast majority of photoreceptors), gene therapies at advanced disease stages were as effective as in early stages. Furthermore, the presence of unrescued, dying photoreceptors did not cause the death of rescued cells, indicating that low-efficiency gene therapies still have therapeutic values (25). Although these studies provide proof-of-concept evidence for the long-term efficacy of suboptimal and late stage gene therapies, it remains to be determined whether the above conclusions are generalizable to other retinal degenerations.

Bardet-Biedl syndrome (BBS) is a genetically heterogeneous, pleiotropic disease associated with ciliary dysfunction (27-29). As in many ciliopathies, retinal degeneration is a major phenotypic component of BBS. BBS patients typically present with night blindness during the first decade in their life and become legally blind by the end of their second decade. Although the retinal dystrophy in BBS is often described as RP or rod-cone dystrophy (i.e. primarily a disease in rods), deficits in cone functions are also evident in young patients. Recent studies in BBS mouse models showed that aberrant accumulation of inner segment (IS) proteins in the OS is a primary cause of photoreceptor degeneration and is accompanied by disorganization of the OS structure (30-32).

In the present study, we determined the long-term efficacy of gene therapy and its therapeutic time window in a preclinical model of ciliopathy, BBS type 17. BBS17 is caused by mutations in the Leucine zipper transcription factor-like 1 (LZTFL1) gene on chromosome 3 (33, 34). To achieve the optimal treatment effect at multiple disease stages, we employed a BBS17 mouse model with a removable gene-trap cassette (30) and systemically restored endogenous *Lztfl1* expression at various ages. The effects of gene therapies were evaluated over a period of 6 months after treatment and compared with the natural history of disease.

## RESULTS

### *Lztfl1* gene-trap mice for temporally controlled, optimal treatment

To achieve the optimal treatment effect, we took a genetic rescue approach and used a BBS17 mouse model with a “knockout-first” allele (*Lztfl1*^*gt*^; (30)) (**Figure 1A**). In this allele, the gene-trap that suppresses *Lztfl1* expression can be eliminated by FLP/FRT-mediated site-specific recombination (35-38). Removal of the gene-trap converts the mutant allele to a functional wild-type, and the resulting *Lztfl1* expression is driven by its natural promoter. For temporally controlled systemic rescue, we crossbred *Lztfl1*^*gt/gt*^ mice with *R26*^*FlpoER*^ mice (39), which express tamoxifen (TMX)-inducible FLP recombinase (FlpER) ubiquitously.

**Figure 1.**
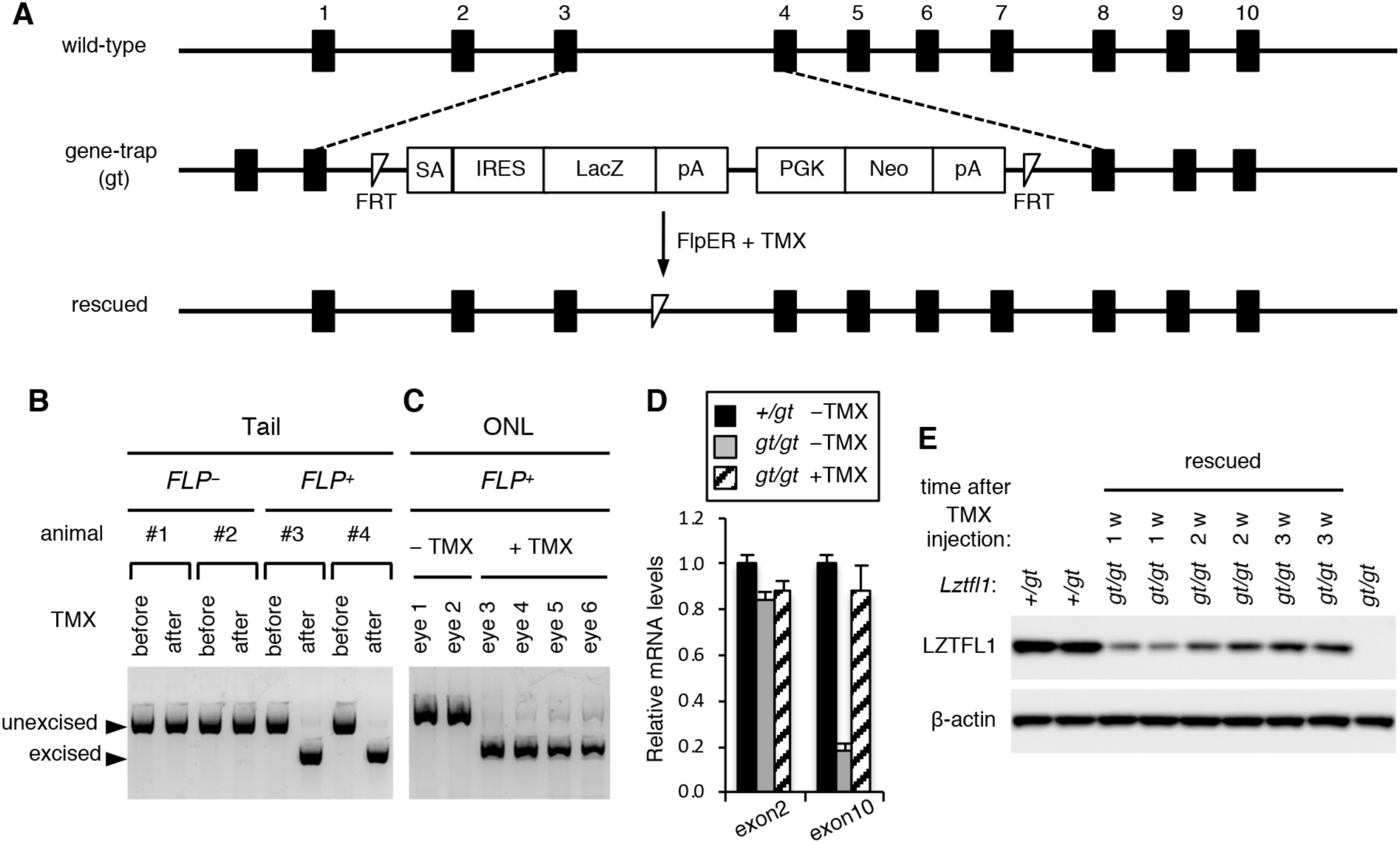
Restoration of *Lztfl1* expression in *Lztfl1*^*gt/gt*^ mice by tamoxifen (TMX)-inducible FLP recombinase. A) Schematic of the wild-type, gene-trap (gt) mutant, and rescued alleles of *Lztfl1*. Black boxes represent *Lztfl1* exons. Exon numbers are described above the black boxes. TMX administration induces excision of the gene-trap by FLP recombinase and converts the gene-trap allele into a functionally normal wild-type allele. SA: splice acceptor site, IRES: internal ribosome entry site, LacZ: β-galactosidase reporter, pA: poly(A) signal, PGK: mouse phosphoglycerate kinase 1 promoter, Neo: neomycin resistance gene. B-C) Validation of the gene-trap cassette excision after TMX injection (at 1 week post-TMX injection). (B) Genomic DNAs from mouse tail snips were isolated before and after TMX injection, and excision of the gene-trap was assessed by PCR. (C) Genomic DNAs were extracted from the outer nuclear layer (ONL) of retinal sections by laser-capture microdissection, and excision of the gene-trap was analyzed by PCR. All mice used are *Lztfl1*^*gt/gt*^, and *FlpER* genotypes are indicated (*FLP*− or *FLP*^+^). D) Restoration of *Lztfl1* expression at the mRNA level (at 1 week post-TMX injection). Total RNA was extracted from the whole eye with or without TMX injection, and the amount of *Lztfl1* mRNA was assessed by qRT-PCR assays. Primers specific to *Lztfl1* exons 2 and 10 were used (n=3 for each group). Error bars represent the standard error of the mean. E) Restoration of *Lztfl1* expression at the protein level. *Lztfl1*^*gt/gt*^;*FLP*^+^ mice were injected with TMX at P30-34, and eyes were collected 1 week (w), 2 weeks, and 3 weeks later for immunoblotting. *Lztfl1*^+*/gt*^;*FLP*^+^ and uninjected *Lztfl1*^*gt/gt*^;*FLP*^+^ mice were used as wild-type and mutant controls, respectively. Each lane represents an individual animal. β-actin was used as a loading control.

We first tested whether the gene-trap cassette could be eliminated by TMX injection as intended. To this end, 1-month old *Lztfl1*^*gt/gt*^ mice with and without FlpER were injected with TMX, and tail snips were collected before and 1 week after TMX injection (see the Materials and Methods section for TMX administration regimen). PCR analyses of the tail snip DNAs showed efficient excision of the gene-trap upon TMX administration in *FLP*^+^ but not in *FLP*− mice (**Figure 1B**). For more direct evaluation of the excision in photoreceptors, the outer nuclear layer (ONL) of the retina was isolated by laser-capture microdissection, and genomic DNAs were extracted and analyzed by the same PCR analysis. As shown in **Figure 1C**, TMX administration induced efficient excision of the gene-trap in the photoreceptor cell layer in TMX-injected *Lztfl1*^*gt/gt*^;*FLP*^+^ mice. In contrast, excision events were not detected without TMX, indicating that FlpER activities are negligible in the absence of TMX (**Figure 1B and C**).

We then measured *Lztfl1* mRNA levels in the eye by quantitative reverse transcription PCR (qRT-PCR) (**Figure 1D**). *Lztfl1* exon 2 is upstream of the gene-trap and transcribed in both wild-type and gene-trap alleles, whereas exon 10 is downstream and expected to be not transcribed in the gene-trap allele because of the presence of two transcription termination signals within the gene-trap (**Figure 1A**). Consistent with this, we found that *Lztfl1* mRNAs containing exon 10 were markedly reduced in *Lztfl1*^*gt/gt*^;*FLP*^+^ mice compared with that of *Lztfl1*^+*/gt*^;*FLP*^+^ animals. The residual amount of *Lztfl1* exon 10 detected in *Lztfl1*^*gt/gt*^;*FLP*^+^ mice is likely derived from read-through transcription. To assess *Lztfl1* expression after treatment, eyes were collected from *Lztfl1*^*gt/gt*^;*FLP*^+^ mice 1 week post-TMX injection (PTI). As shown in **Figure 1D**, *Lztfl1* exon 10 levels were restored to that of *Lztfl1*^+*/gt*^ mice after treatment. Exon 2 levels remained unchanged regardless of the treatment. Absence of sequence alterations in re-expressed *Lztfl1* mRNAs was confirmed by sequencing *Lztfl1* cDNAs (from exon 2 to exon 8) (**Figure S1**).

Finally, we validated restoration of *Lztfl1* expression at the protein level (**Figure 1E**). One-month old *Lztfl1*^+*/gt*^;*FLP*^+^ and *Lztfl1*^*gt/gt*^;*FLP*^+^ mice were treated with TMX, and eyes were collected 1, 2, and 3 weeks PTI. LZTFL1 proteins were readily detectable by immunoblotting in *Lztfl1*^+*/gt*^;*FLP*^+^ and TMX-injected *Lztfl1*^*gt/gt*^;*FLP*^+^ animals at all time points tested, but not in uninjected mutant mice. In addition, the migration rate of LZTFL1 from rescued animals was identical to that from normal controls in SDS-PAGE, indicating that re-expressed proteins were full-length. Interestingly, despite the near immediate restoration of *Lztfl1* expression at the mRNA level (i.e. within 1 week after TMX injection; **Figure 1D**), restoration at the protein level was significantly delayed: LZTFL1 protein levels were relatively low at 1 week PTI and gradually increased until 3 weeks PTI. In summary, our data demonstrate that the gene-trap cassette that blocks *Lztfl1* expression is efficiently removed by FLP recombinase upon TMX injection and that normal expression of *Lztfl1* can be restored inducibly in our animal model.

### Natural history of retinal degeneration in *Lztfl1*^*gt/gt*^ mice

To evaluate the efficacy of gene therapy at multiple disease stages, it is crucial to determine the natural history of disease and use it as a reference point. Therefore, we established the natural history of retinal degeneration in our mouse model.

We first assessed photoreceptor cell loss with age. To this end, we collected serial sections from the central retina and counted the number of rows of photoreceptor cell nuclei stained with 4′,6-Diamidine-2′-phenylindole dihydrochloride (DAPI) (**Figure 2A-C).** The photoreceptor cell layer progressively became thinner in *Lztfl1*^*gt/gt*^ mice. Until post-natal (P) day P21, loss of photoreceptors was marginal (∼10% loss on average; n=4). Thereafter, photoreceptors steadily degenerated and by 6 months of age only 1 or 2 rows of nuclei were left (∼91% loss; n=4). Since the vast majority of photoreceptors (∼97%) in mouse retinas are rods, reduction of the photoreceptor cell layer is mostly accounted for by the loss of rods. To assess the loss of cones, we examined photoreceptor cells expressing *Gnat2*, cone-specific transducin α, in *Lztfl1*^*gt/gt*^ mice. Cone degeneration was noticeable at P21 and steadily progressed with age (**Figure 2B**). The rate of cone degeneration was proportionate to overall loss of the photoreceptor cell layer. In addition, cone OSs appeared to be short even at P21, suggesting that cone OS assembly is impaired in *Lztfl1* mutant retinas (**Figures 2B** and **S2**). As previously described (30), mislocalization of cone opsins (OPN1MW) was also observed in *Lztfl1*^*gt/gt*^ mice (**Figure S2**). These data indicate that both rods and cones are affected by the loss of LZTFL1.

**Figure 2.**
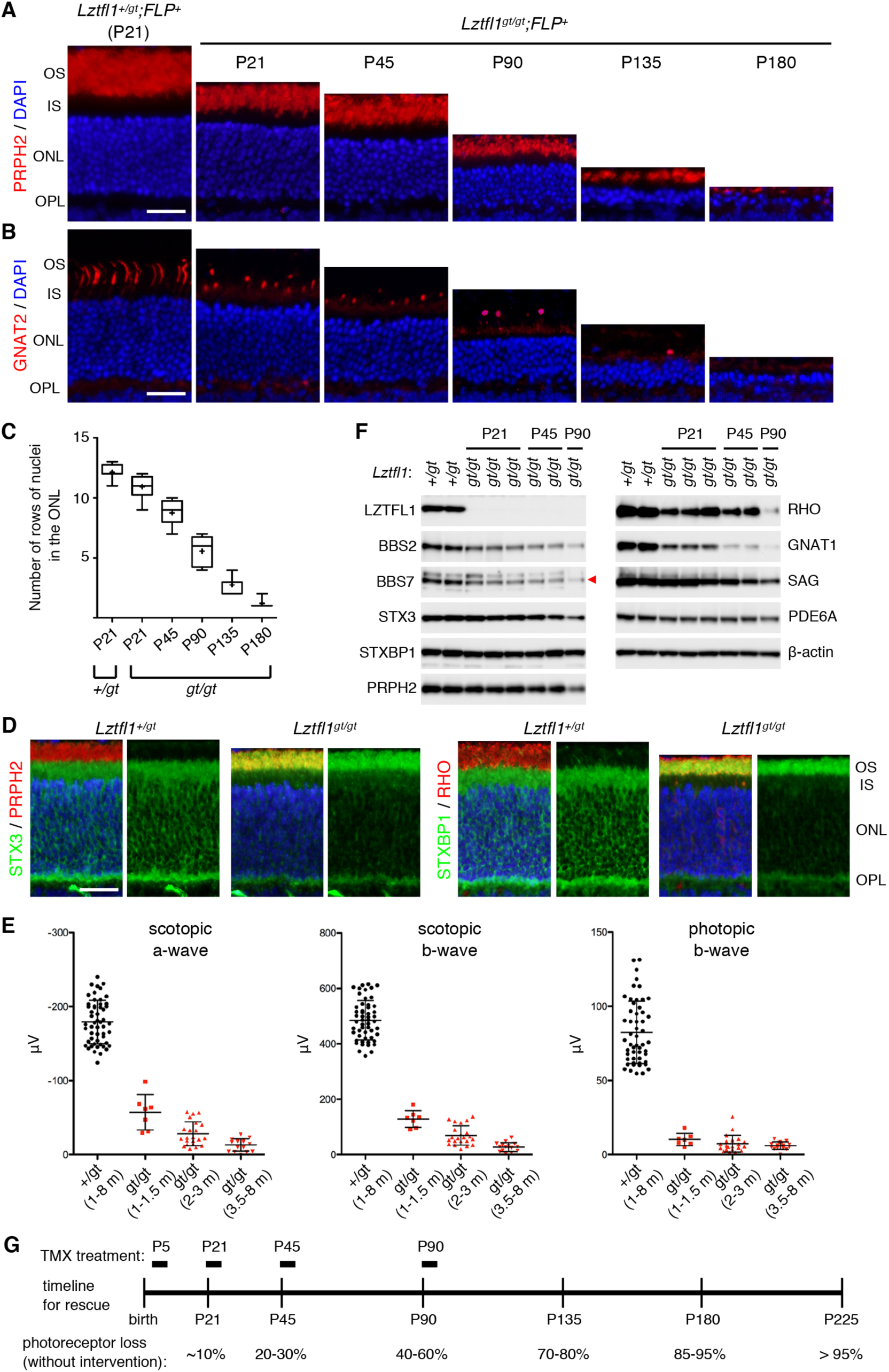
Natural history of photoreceptor degeneration in *Lztfl1*^*gt/gt*^ mice. A) Progressive loss of photoreceptors in *Lztfl1*^*gt/gt*^ mice. Retinal sections from *Lztfl1*^+*/gt*^;*FLP*^+^ and *Lztfl1*^*gt/gt;*^*FLP*^+^ mice at the indicated ages were stained with anti-PRPH2 antibodies (red) and DAPI (blue; for nuclei). OS: outer segment; IS: inner segment; ONL: outer nuclear layer; OPL: outer plexiform layer. Scale bar denotes 25 μm. B) Loss of cones in *Lztfl1*^*gt/gt*^ mice. Retinal sections were stained with anti-GNAT2 (red) antibodies. Others are same as in (A). C) Quantitation of the photoreceptor cell loss. Serial sections perpendicular to the retinal plane were collected from the central retina, and the number of rows of photoreceptor cell nuclei was counted (n=4 per group; 3 sections per mouse; 2 locations per section (within 250 μm from the center)). Data are shown in a box-and-whisker plot. +: mean, error bars: minimum and maximum. D) Mislocalization of STX3 and STXBP1 to the OS in *Lztfl1*^*gt/gt*^ retinas at P13. Retinal sections from *Lztfl1*^+*/gt*^ and *Lztfl1*^*gt/gt*^ mice were decorated with anti-STX3 (green) and anti-STXBP1 (green) antibodies. Anti-PRPH2 (red) and anti-RHO (red) antibodies were used to delineate the OS. Merged images are shown on the left. DAPI (blue) was used to counterstain the nuclei. Scale bar denotes 25 μm. E) Impaired photoresponse in *Lztfl1*^*gt/gt*^ mice. Scotopic ERG a- and b-wave and photopic ERG b-wave amplitudes are shown in scatter plots. Each point represents the average value of two eyes from individual animals. Error bars represent standard deviation (SD) (n=7-51). F) Immunoblot analysis of whole eye protein extracts from *Lztfl1*^+*/gt*^ and *Lztfl1*^*gt/gt*^ mice. Eyes were collected at the indicated ages, and 50 μg of proteins were loaded in each lane after normalization to total protein quantities. Each lane represents an individual animal. β-actin was used as a loading control. BBS7 band was marked with a red arrowhead. G) Timeline of the rescue experiment. TMX was injected at post-natal (P) days P5/8/12 (P5 group), P21- 25 (P21 group), P45-49 (P45 group), and P90-94 (P90 group) to restore *Lztfl1* expression. Natural history of photoreceptor cell loss in *Lztfl1*^*gt/gt*^ mice is summarized at the bottom.

Accumulation of IS proteins STX3 and STXBP1 in the OS has been described in all BBS mouse models examined and in *Cep290* mutant mice (30-32, 40). Mislocalization of these proteins was evident as early as at P13 (the youngest age examined) in *Lztfl1*^*gt/gt*^ mice and observed throughout the length of the OS (**Figure 2D**), indicating that the pathogenic process begins before or when the OS begins to elongate.

We next examined electrical responses of the retina to light by electroretinography (ERG). *Lztfl1*^*gt/gt*^ mice were divided into 3 age groups (1-1.5 months, 2-3 months, and 3.5-8 months old), and scotopic (dark-adapted) ERG a- and b-waves and photopic (light-adapted) ERG b-wave were measured (**Figure 2E**). Control animals (*Lztfl1*^+*/gt*^; 1-8 months of age) were grouped into a single cohort as no age-dependent changes were observed in our colony. In all three age groups and regardless of the light conditions, the capacity of photoreceptors to respond to light was severely diminished in *Lztfl1*^*gt/gt*^ mice compared with *Lztfl1*^+*/gt*^ animals (one-way ANOVA followed by Tukey’s multiple comparison test; *p*<0.001), indicating that both rod and cone functions were severely impaired even in young animals. Also, in scotopic conditions in which rod functions were mainly measured, there was a trend of gradual reduction in both a- and b-wave amplitudes (1-1.5 m vs. 3.5-8 m; *p*<0.001).

To gain more insight into the pathophysiology of impaired visual functions and retinal degeneration, we examined proteins involved in the phototransduction cascade and the OS morphogenesis as well as components of the BBSome (a protein complex composed of eight BBS proteins (41, 42)) by immunoblotting (**Figure 2F**). BBSome subunits, BBS2 and BBS7, showed a moderate (50-60%) reduction at P21 and continued to decrease as photoreceptors degenerated. To test whether loss of LZTFL1 affects BBSome assembly in the retina, protein extracts from normal and *Lztfl1*^*gt/gt*^ eyes were loaded on a 10-40% sucrose gradient and sedimentation rates of individual BBSome components were analyzed by sucrose gradient ultracentrifugation and immunoblotting. As shown in **Figure S3**, all BBSome components in both normal and *Lztfl1*^*gt/gt*^ eyes sedimented together at the same rate. This result is consistent with our prior finding in cultured mammalian cells (43) and indicates that BBSome assembly is not affected by the loss of LZTFL1 despite overall reduction in BBSome quantity. Among the proteins tested, transducin α (GNAT1) was the most sensitive protein to the loss of LZTFL1 and showed ∼80% reduction at P21 and further decreased as photoreceptors degenerated (**Figure 2F**). Rhodopsin (RHO) was also sensitive to the loss of LZTFL1 and showed ∼60-70% reduction at P21. Paucity of these phototransduction proteins is likely to contribute to the severely impaired visual functions in *Lztfl1*^*gt/gt*^ mice before photoreceptor loss. In contrast, visual arrestin (SAG), PRPH2, PDE6A, and STX3 levels were only marginally reduced at P21, and their reduction was roughly proportionate to the loss of photoreceptors. STXBP1 is expressed in not only photoreceptors but also other neurons in the inner retina, and its protein level remained unchanged despite loss of photoreceptors (due to the normalization of protein extracts to total protein quantities before SDS-PAGE). β-actin was used as a loading control and also showed no changes.

Based on the degree of photoreceptor cell loss, we selected 4 ages that represent i) before or very early stage of degeneration (P5), ii) early stage (P21; ∼10% cell loss), iii) mid stage (P45; 20-30% loss), and iv) late stage (P90; 40-60% loss) (**Figure 2G**). *Lztfl1*^*gt/gt*^ mice with FlpER were divided into these 4 age groups, injected with TMX, and the long-term efficacies of early-to-late stage gene therapies were examined. Of note, no differences were observed between TMX-injected and uninjected *Lztfl1*^+*/gt*^;*FLP*^+^ mice with regard to the experimental measurements described in this study: ERG, protein quantity, and protein localization. Therefore, both groups of animals were used as normal controls. Also, TMX-uninjected *Lztfl1*^*gt/gt*^;*FLP*^+^ and TMX-injected *Lztfl1*^*gt/gt*^;*FLP*− mice were indistinguishable, and both groups of animals were used as untreated mutant controls throughout the study. To maximize the gene-trap excision efficiency, all *Lztfl1*^*gt/gt*^ mice used for rescue experiments were homozygous for the *FLP* transgene.

### Restoration of *Lztfl1* expression at very early degeneration stages preserves vision and delays retinal degeneration but does not permanently arrest disease progression

To test the efficacy of gene therapy before degeneration begins, we initially administered TMX to *Lztfl1*^*gt/gt*^;*FLP*^+^ mice between P1 and P7. Our initial administration regimen was a one-time injection of 0.1-0.7 mg of TMX between P1 and P7 (e.g. 0.4 mg at P4 or 0.7 mg at P7). We also tested two-time injections in two nonconsecutive days (e.g. 0.4 mg at P4 followed by 0.7 mg at P7). However, these attempts were not successful because of the toxicity of TMX at “high” doses in neonates, while doses that were not lethal failed to induce more than 30% of excision as determined by tail DNA PCR. Like other BBS mutants (44-49), *Lztfl1*^*gt/gt*^ newborn pups were smaller than their normal littermates and more susceptible to TMX. The highest dose that could achieve more than 80% excision with less than 50% lethality was triple injection of TMX at P5, P8, and P12 (0.5 mg at P5 followed by 0.8 mg at P8 and 1 mg at P12). Therefore, we used this dose and age (the P5 group in **Figure 2G**) to examine whether the aforementioned molecular phenotypes and photoreceptor degeneration could be rescued by gene therapy in *Lztfl1* mutant mice.

We first confirmed restoration of *Lztfl1* expression at the protein level in P5-rescued *Lztfl1* mutants. At 1 month PTI, LZTFL1 proteins were readily detectable in treated *Lztfl1*^*gt/gt*^;*FLP*^+^ retinas but not in untreated retinas by immunoblotting (**Figure 3A**). We also examined restoration of other proteins that showed significant reduction in *Lztfl1* mutants. Although not prominent at this age, restoration of *Lztfl1* expression resulted in a slight increase (5-30%) of BBS2 and BBS7 in treated retinas. GNAT1, which was highly sensitive to the loss of LZTFL1 (**Figure 2F**), was markedly increased (∼2.5-fold) in treated retinas (**Figure 3A**). At 3 months and 6 months PTI, *Lztfl1* expression was maintained and BBS2, BBS7, RHO, GNAT1, and STX3 levels were well preserved in treated animals (**Figure 3B and C**). Taken together, these data demonstrate that restoration of *Lztfl1* expression between P5 and P12 effectively prevents degradation of phototransduction proteins and BBSome components for more than 6 months.

**Figure 3.**
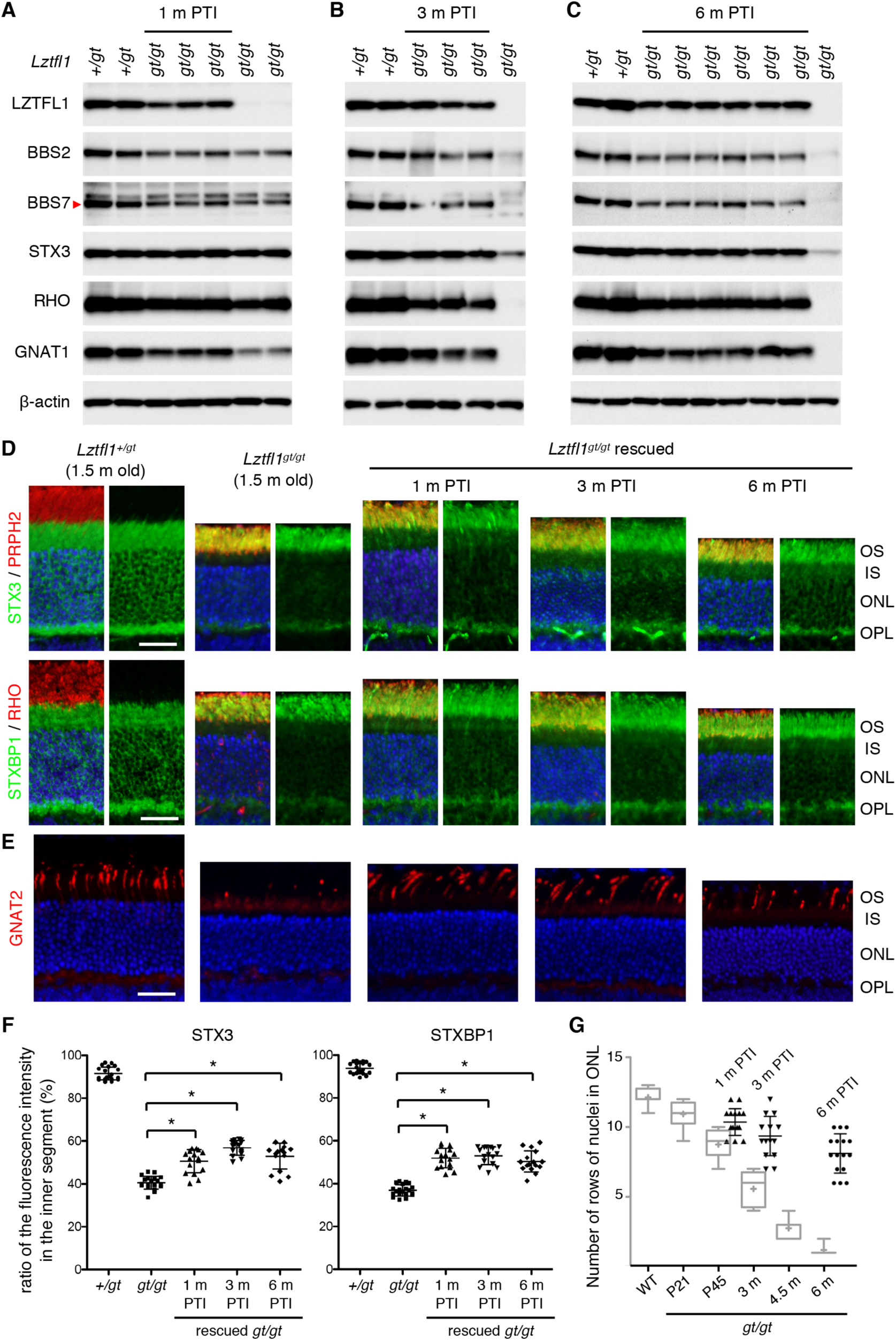
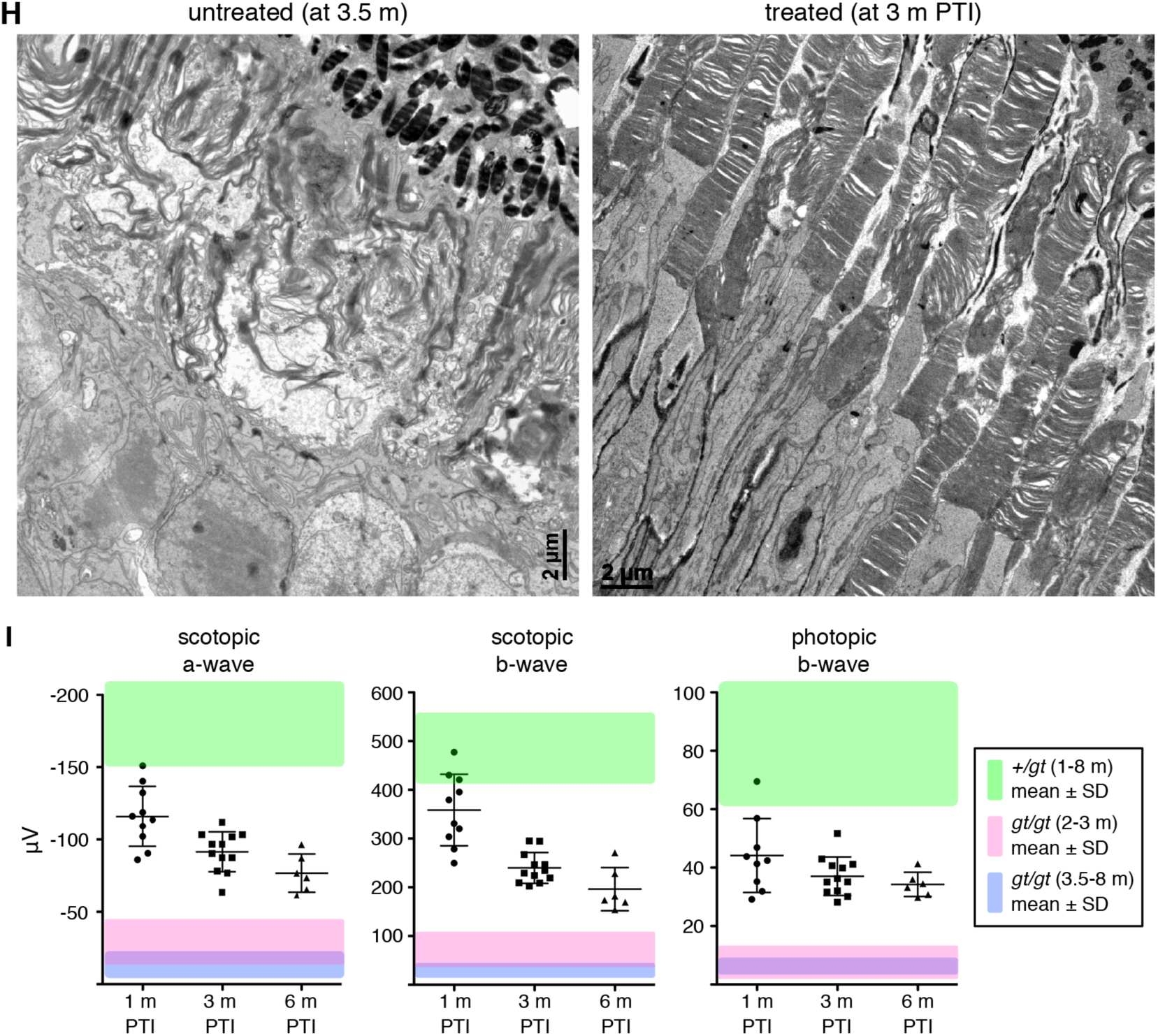
Restoring *Lztfl1* expression at P5/8/12 has long-term beneficial effects but does not permanently halt retinal degeneration. A-C) Preservation of BBSome components and phototransduction proteins at 1 month (A), 3 months (B), and 6 months (C) post-TMX injection (PTI). Age-matched *Lztfl1*^+*/gt*^ and *Lztfl1*^*gt/gt*^ mice (unrescued) were used as positive and negative controls. Each lane represents an individual mouse. D) Retinal degeneration is significantly delayed but protein mislocalization is only partly corrected in the P5-rescue group. Others are same as in Figure 2D. Scale bar denotes 25 μm. E) Long-term preservation of cones in the P5-rescue group. Anti-GNAT2 antibody (red) was used to visualize cone OSs. F) Quantitation of STX3 and STXBP1 localization in the inner segment. Depicted are the ratio of integrated fluorescence intensities in the inner segment relative to the photoreceptor cell layer. Data are from 4-5 mice (2 sections per mouse and 2 areas per section) and mean ± SD (error bar) is marked. Asterisks indicate statistical significance (one-way ANOVA followed by Tukey’s multiple comparison test; *p*<0.01). G) Quantitation of the photoreceptor cell layer in treated animals. The number of rows of photoreceptor cell nuclei was counted in treated animals. For direct comparison with unrescued *Lztfl1*^*gt/gt*^ mice, data were presented in a scatter plot and overlaid on the graph shown in Figure 2C (light grey). Mean ± SD is shown (n=4; 2 sections per mouse; 2 places per section). H) Transmission electron micrographs (TEM) of the retinas from untreated (left; 3.5-months old) and treated (right; at 3 months PTI) *Lztfl1*^*gt/gt*^;*FLP*^+^ mice. I) ERG responses of *Lztfl1*^*gt/gt*^;*FLP*^+^ mice treated at P5/8/12. Scotopic ERG a- and b-wave and photopic ERG b-wave amplitudes are shown in scatter plots. Error bars represent SD. Green-shaded area denotes mean ± SD of normal mice (1-8 months old) from Figure 2E. Pink-and blue-shaded areas mark mean ± SD of untreated *Lztfl1*^*gt/gt*^ mice (2-3 months old and 3.5-8 months old, respectively) from Figure 2E.

We then examined photoreceptor cell loss and the localization of STX3 and STXBP1. Since the OS begins to develop at P5 in mice (50, 51) and is essentially absent or rudimentary at the time of treatment, we used 1.5-month old animals with fully differentiated retinas as a reference point for the localization study. In normal retinas, STX3 and STXBP1 are almost exclusively found in the IS (hereafter our use of the term IS encompasses all parts of photoreceptors including soma, axon, and synaptic terminal but excluding the OS) (**Figure 3D**). In 1.5-month old *Lztfl1*^*gt/gt*^ mice, however, only 40% of STX3 and STXBP1 were found in the IS and the majority (∼60%) localized to the OS (**Figure 3D and F**). Most of the IS pool remaining was localized to the synaptic terminal but the density there was significantly reduced compared to normal retinas. Thinning of the ONL was also apparent in *Lztfl1*^*gt/gt*^ mice. At 1 month PTI, the ONL of treated *Lztfl1*^*gt/gt*^;*FLP*^+^ mice was about 2 rows of nuclei thicker than that of untreated mice (n=4) (**Figure 3G**). Preservation of photoreceptors was more conspicuous at 3 months and 6 months PTI, demonstrating the long-term efficacy of early stage gene therapy. The ONL, however, became gradually thinner as treated animals aged, suggesting that retinal degeneration continued at a slower rate. Mislocalization of STX3 and STXBP1 was partly corrected (**Figure 3D and F**). The ratio of these proteins in the IS was significantly increased in treated animals (one-way ANOVA followed by Tukey’s multiple comparison test; *p*<0.01). To our surprise, however, 40-60% of STX3 and STXBP1 still mislocalized to the OS. This mislocalization phenotype was not corrected until 6 months PTI, when >90% of photoreceptors were lost in untreated retinas.

We next examined the survival and morphology of cones (**Figure 3E**). At 1 month PTI, significantly more cones were found in treated retinas compared with age-matched, untreated retinas. In addition, the length of the cone OS was comparable to that of normal cones. These data indicate that restoration of *Lztfl1* expression not only prevents cell death but also restores OS formation in cones. However, similar to the slow but continuous loss of rods, the number of cones gradually decreased with age.

Ablation of BBS gene function causes disc morphogenesis defects and disorganization of the OS ultrastructure (30-32). We examined whether restoring *Lztfl1* expression at P5-12 could preserve the OS ultrastructure, using transmission electron microscopy (TEM) (**Figures 3H** and **S4**). In 3.5-month old *Lztfl1*^*gt/gt*^ mouse retinas, individual OSs could not be distinguished. The space between the pigmented epithelium and the photoreceptor IS was filled with elongated and disorganized disc membranes and various sizes of extracellular vesicular structures. At 3 months PTI, the OS ultrastructure of treated *Lztfl1*^*gt/gt*^ mice was significantly better preserved compared with untreated animals. In many cells, discs were densely stacked as in normal photoreceptors. However, a subset of cells had discs aligned along the longitudinal axis of the cell, indicating that disruption of the disc morphogenesis process was still occurring.

Finally, ERG responses of the retina were measured (**Figure 3I**). At 1 month PTI, scotopic ERG a- and b-waves were remarkably better in treated animals than in the untreated ones (one-way ANOVA followed by Tukey’s multiple comparison test; *p*<0.01). A few animals’ ERG amplitudes were within or close to the mean ± SD range of normal animals (green-shaded area). Photopic ERG b-wave amplitudes were also significantly better (*p*<0.01). At 3 months and 6 months PTI, all ERG responses remained more robust than age-matched untreated animals (pink- and blue-shaded areas; *p*<0.01). However, consistent with the continuous loss of photoreceptor cells, ERG amplitudes gradually decreased in treated *Lztfl1*^*gt/gt*^;*FLP*^+^ animals. Of note, decreases in ERG responses occurred in all animals, including the ones with near-normal ERG amplitudes at 1 month PTI. We kept 3 treated animals until 9 months PTI and examined protein mislocalization, ONL thickness, and ERG responses. In these animals, ERG amplitudes and ONL thickness decreased further while protein mislocalization was unchanged (**Figure S5**).

In summary, our data indicate that restoration of *Lztfl1* expression at a very early stage of degeneration has long-term therapeutic effects (preservation of phototransduction proteins and visual functions and long-term survival of photoreceptor cells), but it does not completely correct protein mislocalization nor permanently stop degeneration.

### The efficacy of gene therapy decreases in *Lztfl1* mutant mice as gene expression is restored at later disease stages

We then examined the efficacy of gene therapies at later disease stages. To this end, TMX was administered at 3 different ages (see **Figure 2G**). The P21 group was injected at P21-25 (5 consecutive days of injection; 1 injection/day). The P45 and P90 groups were injected at P45-49 and P90-94, respectively (5 consecutive days of injection; 1 injection/day). As in the P5 group, protein localization and expression, photoreceptor cell loss, and retinal functions were examined at 1 month, 3 months, and 6 months PTI.

Administration of TMX at P21-25 restored *Lztfl1* expression in *Lztfl1*^*gt/gt*^;*FLP*^+^ mice and its expression was maintained for more than 6 months (the last time point examined) (**Figure 4A-C**). Restoration of *Lztfl1* expression at this age prevented loss of the BBSome components and the phototransduction proteins for at least 3 months PTI. However, this rescue effect diminished by 6 months PTI despite continued expression of *Lztfl1*, suggesting resumption or continuation of retinal degeneration. We then assessed photoreceptor cell loss and protein localization by immunohistochemical studies. In all ages examined, the photoreceptor cell layer of treated mice was significantly thicker than that of age-matched, untreated animals (compare **Figure 4D** and **Figure 2A**). Cone cells were also preserved, and their OSs were elongated (**Figure 4E**). However, consistent with the biochemical data in **Figure 4A-C**, photoreceptor degeneration continued and the photoreceptor cell layer became thinner with age. This indicates that gene therapy at P21-25 delays photoreceptor cell loss in *Lztfl1* mutant mice but does not permanently prevent it. In addition, despite some improvement in distribution, STX3 and STXBP1 mislocalization was not fully corrected in P21-rescued eyes in all ages examined (**Figure 4D**). Finally, ERG data showed that both rod and cone functions were significantly better than untreated animals initially but declined over time (**Figure 4F**). At 1 month PTI, scotopic ERG a- and b-waves and photopic b-wave were significantly higher than those of untreated mice at similar ages (pink-shaded area; *p*<0.01). These ERG amplitudes, however, gradually diminished with age. The rate of ERG decline in this group was faster than that of the P5 group. Overall, our data indicate that gene therapy at P21-25 in *Lztfl1* mutant mice delays retinal degeneration but it does not permanently halt it.

**Figure 4.**
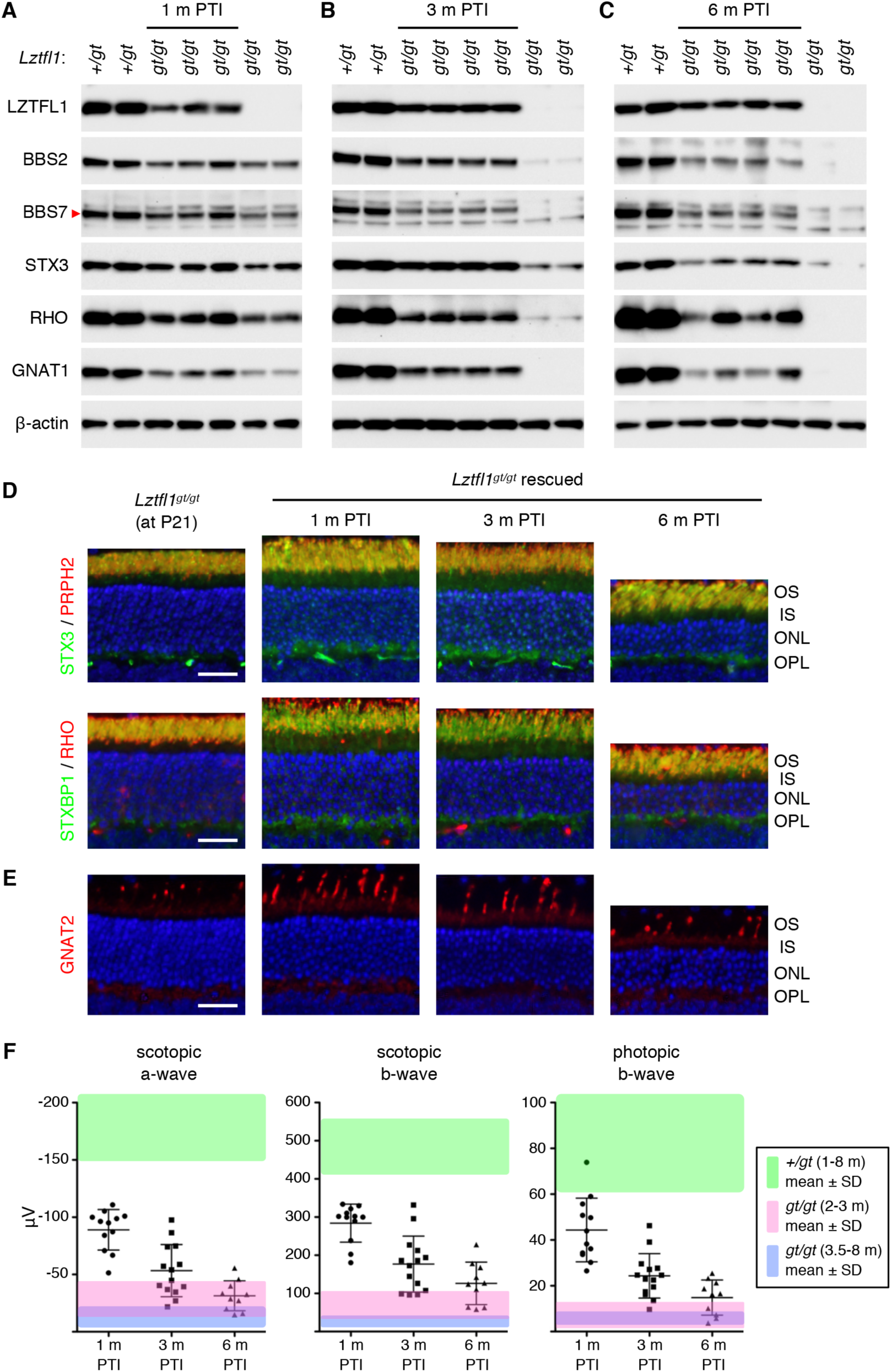
Long-term effects of *Lztfl1* gene therapy in the P21-rescue group. A-C) Immunoblot analyses of P21-treated *Lztfl1*^*gt/gt*^;*FLP*^+^ mice at 1 month (A), 3 months (B), and 6 months (C) PTI. D) STX3 and STXBP1 localization in P21-treated *Lztfl1*^*gt/gt*^;*FLP*^+^ mice. Retinal sections from untreated *Lztfl1*^*gt/gt*^ mice at P21 are shown on the left as a reference point. Scale bar: 25 μm. E) Survival of cones in P21-treated *Lztfl1*^*gt/gt*^;*FLP*^+^ mice. F) ERG responses of *Lztfl1*^*gt/gt*^;*FLP*^+^ mice treated at P21-25. Others are same as in Figure 3.

We next investigated the efficacy of gene therapy at a mid-disease stage (20-30% cell loss). *Lztfl1*^*gt/gt*^;*FLP*^+^ mice were administered with TMX at P45-49 and the efficacy of gene therapy was assessed as above. Restoration of *Lztfl1* expression was confirmed by immunoblotting at all time points examined (**Figure 5A-C**). However, BBS2, BBS7, RHO, and GNAT1 protein levels were only marginally higher in treated animals compared with age-matched, untreated animals at 1 month and 3 months PTI. At 6 months PTI, these proteins were barely detectable in treated animals. Immunolocalization and histological studies indicated that protein mislocalization phenotype was not corrected and that photoreceptor degeneration continued in treated animals, despite some delay compared with untreated mice (**Figure 5D** and **E**). Cone OSs appeared to be slightly elongated in a subset of cells at 1 month PTI, but degeneration of cones continued and GNAT2-expressing cones were undetectable at 6 months PTI (**Figure 5E**). To test whether restoration of *Lztfl1* expression at a mid-disease stage has any positive effect on the OS ultrastructure, retinas from treated animals and their untreated littermates were examined by TEM at 1 month PTI. As shown in **Figures 5F** and **S6**, some photoreceptors in the treated retinas had OSs with stacked discs, while no discernable OS structure could be found in untreated retinas. These data indicate that restoring *Lztfl1* expression at a mid-disease stage at least transiently improves the OS ultrastructure. ERG data also showed a transient improvement followed by a decline in visual responses (**Figure 5G**). Although there were statistically significant differences in scotopic ERG b-wave values at all 3 times points and photopic ERG b-wave at 1 month PTI compared with age-matched untreated animals (pink- and blue-shade areas; *p*<0.01), the differences were small and essentially extinct by 6 months PTI. These results indicate that restoration of *Lztfl1* expression at a mid-disease stage has short-term therapeutic effects of only marginal magnitude.

**Figure 5.**
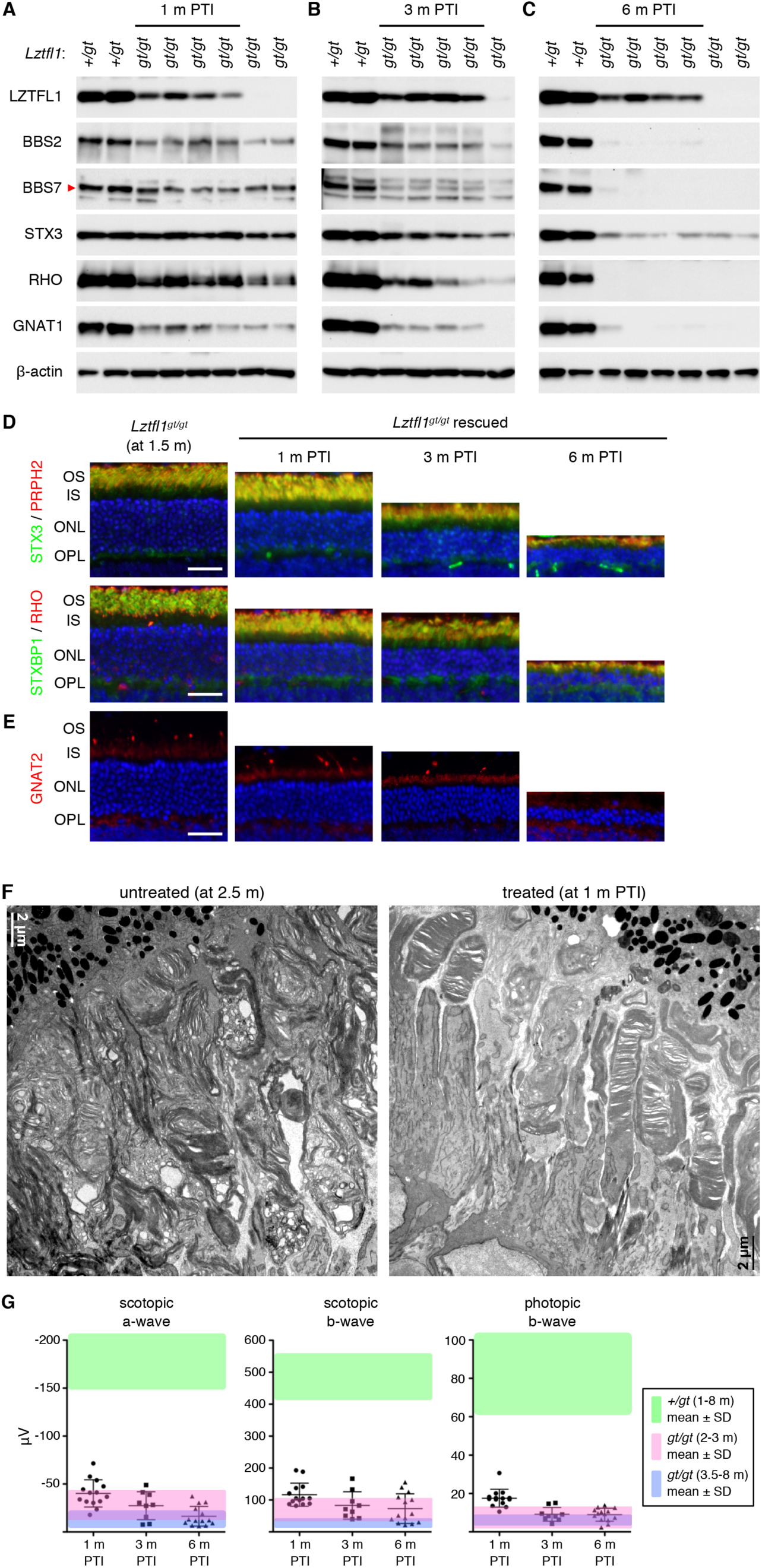
Rescue at a mid-disease stage (P45-49) results in only a temporary and marginal improvement in vision. A-C) Immunoblot analyses of P45-treated *Lztfl1*^*gt/gt*^;*FLP*^+^ mice at 1 month (A), 3 months (B), and 6 months (C) PTI. D) STX3 and STXBP1 localization in P45-treated *Lztfl1*^*gt/gt*^;*FLP*^+^ mice. Retinal sections from 1.5-month old, untreated *Lztfl1*^*gt/gt*^ mice are shown on the left as a reference point. Scale bar: 25 μm. E) Survival of cones in P45-treated *Lztfl1*^*gt/gt*^;*FLP*^+^ mice. F) TEM images of the retinas from untreated (left; 2.5-months old) and treated (right) *Lztfl1*^*gt/gt*^;*FLP*^+^ mice at 1 month PTI. G) ERG responses of *Lztfl1*^*gt/gt*^;*FLP*^+^ mice treated at P45-49. Others are same as in Figure 3.

Finally, we evaluated the efficacy of gene therapy at a late disease stage (at P90-94; ∼40-60% of cell loss). Consistent with the mid-disease stage results, treatment at this age only marginally improved the levels of BBSome components and phototransduction cascade proteins (**Figure 6A** and **B**). Immunolabeling of retinal sections and ERG analyses also indicated that photoreceptors continued to degenerate with little or no detectable therapeutic effect (**Figure 6C-E**). Due to the near complete loss of photoreceptors and lack of rescue effects at the 3 months PTI time point, no further analyses were conducted at later time points. To test whether *Lztfl1* gene therapy is ineffective in general at this age, we examined the fertility of P90-rescued *Lztfl1* mutant males. BBS mutant males including *Lztfl1* mutants are infertile due to a defect in spermatogenesis (44-49). We set up breeder cages with treated and untreated *Lztfl1*^*gt/gt*^;*FLP*^+^ males together with wild-type females (*Lztfl1*^+*/*+^). All treated *Lztfl1* mutant males (n=5) sired offspring (5-8 pups per litter) within 4 months after treatment, whilst untreated males (n=3) produced no pups, indicating that the male infertility phenotype was rescued by restoring *Lztfl1* expression at 3 months of age. Our data indicate that restoring *Lztfl1* expression in the retina at late disease stages has little or no therapeutic effect on retinal degeneration.

**Figure 6.**
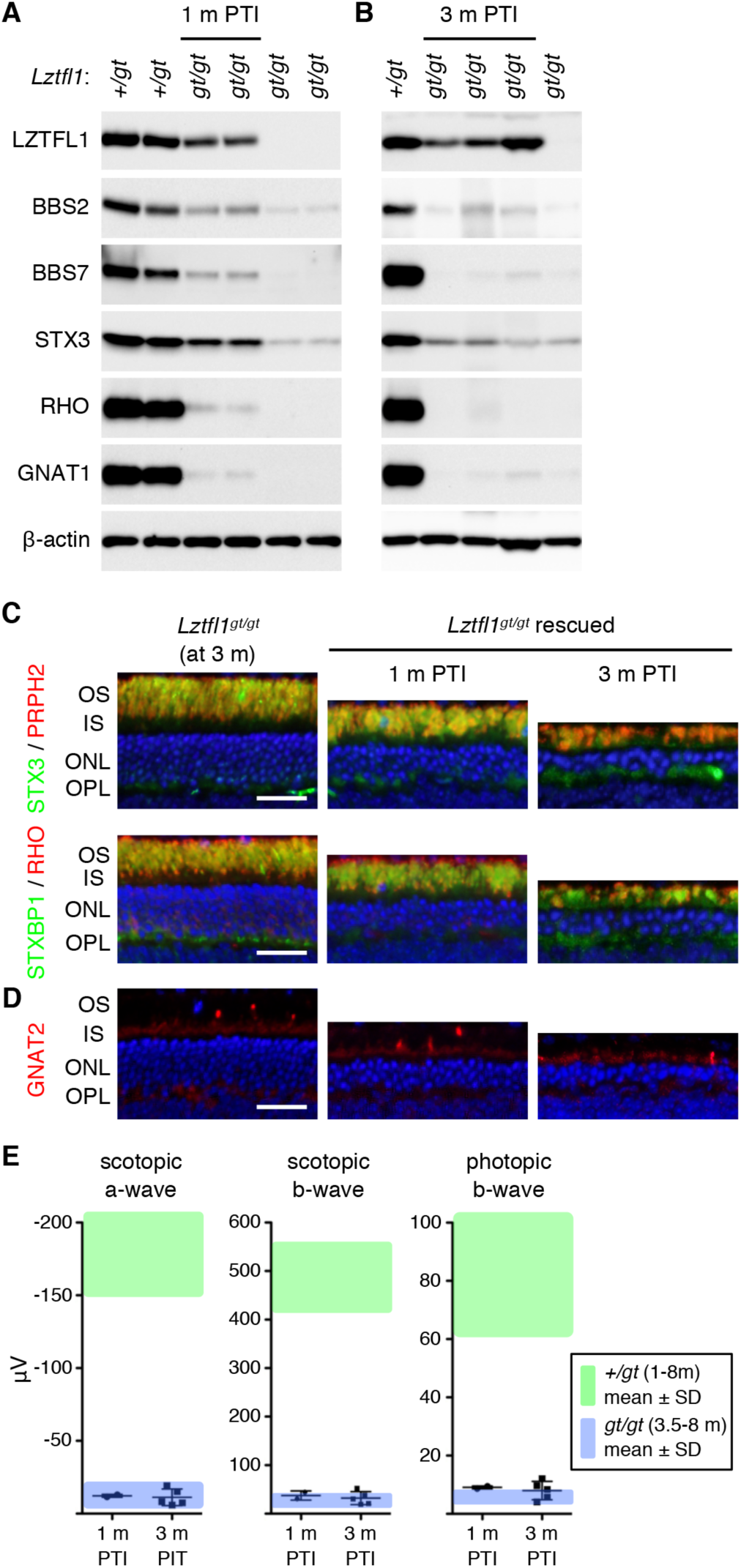
Restoring *Lztfl1* expression at an advanced stage (P90-94) has little therapeutic effect. A-B) Immunoblot analyses of P90-treated *Lztfl1*^*gt/gt*^;*FLP*^+^ mice at 1 month (A) and 3 months (B) PTI. C) STX3 and STXBP1 localization in P90-treated *Lztfl1*^*gt/gt*^;*FLP*^+^ mice. Scale bar: 25 μm. D) Survival of cones in P90-treated *Lztfl1*^*gt/gt*^;*FLP*^+^ mice. E) ERG responses of *Lztfl1*^*gt/gt*^;*FLP*^+^ mice treated at P90-94. Others are same as in Figure 3.

## DISCUSSION

In this work, we show that the efficacy of BBS17 retinal gene therapy depends on the disease stage at which the mutated gene expression is restored. Restoration of *Lztfl1* expression at early stages preserves both rods and cones for a long term and significantly improves visual function. However, the therapeutic efficacy gradually declines as *Lztfl1* expression is restored at later stages, with no treatment effect at the advanced stage despite the presence of a considerable number of photoreceptors (40-60%) at the time of treatment and successful restoration of gene expression. Furthermore, even at early stages, the protein mislocalization phenotype was not fully corrected and retinal degeneration continued at a slower rate. Our results contrast with the previous study by Koch *et al*. (24), which employed a similar genetic rescue approach to achieve the “optimal” treatment effect and found that retinal degeneration caused by mutations in *Pde6b* could be arrested even when less than 40% of photoreceptors were remaining. Our study indicates that at least in certain retinal degenerations there is a limited time window for successful retinal gene therapy.

Multiple lines of evidence indicate that the persistent degeneration in treated *Lztfl1* mutant mice is not due to inefficient excision of the gene-trap cassette. First, PCR analyses of genomic DNAs isolated by laser-capture microdissection of the ONL show that excision of the gene-trap occurred in the vast majority of photoreceptors. Second, qRT-PCR data to measure *Lztfl1* mRNA levels also indicate that the quantity of *Lztfl1* mRNAs in treated animals was comparable to that of normal animals. Third, LZTFL1 proteins were detected in the retina regardless of the ages after treatment. Importantly, multiple phenotypes were rescued or improved after treatment. These include preservation of the phototransduction proteins, prolonged survival of both rods and cones, improved OS morphology, and improved ERG responses. Improved OS morphology observed by TEM strongly argues that at least a subset (probably a majority) of photoreceptors were indeed rescued in treated animals. In addition, *Lztfl1* cDNA sequencing and immunoblotting data indicate that LZTFL1 proteins expressed from the rescued allele were normal full-length proteins. Therefore, although we cannot completely rule out the possibility of non-cell autonomous cell death due to extrinsic toxic factors derived from unrescued dying cells, our data suggest that *Lztfl1* expression was restored in most photoreceptors and nevertheless retinal degeneration continued at a slower rate.

To our surprise, protein mislocalization phenotype was partly but not completely corrected after LZTFL1 restoration. We were not able to find any individual photoreceptors with normal STX3 and STXBP1 localization (i.e. IS-restricted localization) even at 9 months PTI in the P5-rescue group. Essentially all photoreceptors die and disappear by this age in untreated *Lztfl1* mutant mice. These findings suggest that the partial improvement in protein localization on the macroscopic scale is not due to the presence of two populations of photoreceptors (i.e. rescued cells with normal localization patterns and unrescued cells with reversed localization). Instead, our data suggest that protein mislocalization continues to occur after the re-expression of LZTFL1. In addition, normal photoreceptor OSs renew constantly (52), and thus STX3 and STXBP1 mislocalized to the OS before LZTFL1 restoration are expected to be removed by the OS renewal over time. The persistent protein mislocalization suggests that mechanisms that prevent STX3 and STXBP1 mislocalization at the connecting cilium (53) are not fully repaired. Continued protein mislocalization may contribute to the continued retinal degeneration in treated mice.

Persistent protein mislocalization and continuous progression of retinal degeneration in treated *Lztfl1* mutant mice is surprising, because in our prior study (31) restoration of *Bbs8* expression in *Bbs8*^*gt/gt*^ mice between P9 and P15 using the same FLP driver (*R26*^*FlpoER*^) fully rescued the protein mislocalization phenotype and prevented further progression of degeneration. Restoration at P25-29, which is after the terminal differentiation of photoreceptors, also fully corrected protein mislocalization (**Figure S7**). Since we used the same FLP driver, differential outcomes in *Lztfl1*^*gt/gt*^ and *Bbs8*^*gt/gt*^ mice cannot be explained by differences in FLP recombinase expression or activity. In addition, complete rescue in *Bbs8*^*gt/gt*^;*FLP*^+^ mice and the lack of protein mislocalization in TMX-injected *Lztfl1*^+*/gt*^;*FLP*^+^ mice argues against the possibility of FRT-independent activity of FLP as a cause of STX3/STXBP1 mislocalization.

What, then, is responsible for the contrasting outcomes in *Lztfl1*^*gt/gt*^ and *Bbs8*^*gt/gt*^ mice? Although the answer to this question is currently unknown, one potential clue is the difference between expression profiles of *Lztfl1* and *Bbs8* during photoreceptor maturation. In mice, the connecting cilium assembles at P3-5 and the OS begins to elongate afterwards (50, 51). Expression of genes involved in ciliogenesis such as the ones encoding IFT (intraflagellar transport) proteins and CEP290 peaks during the connecting cilium assembly and declines significantly as photoreceptors mature (**Figure S8A** and **B**) (also see Fig. S1 in (31)), implying a high demand of these proteins during the connecting cilium assembly. In contrast, expression of BBSome components begins to increase at P4 and peaks at P13-21, but it does not decline as much as IFT proteins in mature photoreceptors (∼50% decrease from the peak). Expression of *Lztfl1* follows the pattern of IFT genes rather than BBSome components. In addition, in the “photoreceptorless” retinas of *Ift88*^*fl/fl*^;*rhodopsin-Cre* conditional knockout mice, BBSome components are barely detectable while IFT proteins and LZTFL1 are readily detected (**Figure S8C** and **D**). This suggests that BBSome components are mostly expressed in photoreceptors in the retina, but IFT proteins and LZTFL1 are expressed in not only photoreceptors but also other cells. At the molecular level, LZTFL1 links BBSomes to IFT particles such that BBSomes can be transported by IFT trains along the axoneme (54). Therefore, although inactivation of *LZTFL1* causes a ciliopathy that falls into the clinical category of BBS, its expression profile and functions are more closely related to those of IFT genes rather than BBSome components. Gene therapies targeting diseases associated with connecting cilium or ciliogenesis defects might require treatment before the connecting cilium assembly, while other diseases associated with defects at later phases in photoreceptor differentiation may be more amenable to later stage treatment. In this regard, it is noteworthy that early-stage gene therapy of LCA5, which is caused by loss of Lebercilin that interacts with IFT particles and mediates protein trafficking to the OS (55, 56), at P5 and P15 did not arrest retinal degeneration despite improvement in functional vision and delay in degeneration (57).

Another potential factor that may contribute to the differential outcomes in *Lztfl1*^*gt/gt*^ and *Bbs8*^*gt/gt*^ mice is residual activities of the mutant alleles. “Knockout-first” gene-trap alleles are designed to interrupt target genes’ expression by hijacking mRNA splicing events and replacing target genes’ downstream exons with a reporter followed by transcription termination. However, low level expression of target genes is often observed presumably because of read-through transcription and skipping of the gene-trap exon during splicing. Because of this, gene-trap mutant animals often display milder phenotypes than null mutants. Indeed, very small amounts of LZTFL1 and BBS8 proteins are detectable in *Lztfl1*^*gt/gt*^ and *Bbs8*^*gt/gt*^ mice, respectively. However, the quantity of LZTFL1 proteins expressed in *Lztfl1*^*gt/gt*^ mice is less than 0.2% of normal animals (*Lztfl1*^+*/gt*^), while BBS8 expressed in *Bbs8*^*gt/gt*^ mice is 5-6% of wild-type (*Bbs8*^+*/*+^) (31). The rate of retinal degeneration is also slightly slower in *Bbs8*^*gt/gt*^ mice compared with *Bbs8* null (*Bbs8*^-/-^) mice. Interestingly, *Pde6b* mutant mice used to demonstrate efficient rescue at advanced disease stages also possessed hypomorphic alleles (*Pde6b*^*H620Q/Stop*^ compound heterozygotes), and retinal degeneration in this model was significantly slower than in *Pde6b*^*rd1/rd1*^ null mutants (24, 58-60). Residual activities of mutated genes might be important for full recovery of photoreceptor cells when normal gene expression is restored. If this is the case, there might be a correlation between specific mutations that patients have and the long-term efficacy of retinal gene therapy.

Our study demonstrates that there is a “point of no return” (61) in certain inherited retinal diseases. In *Lztfl1* mutant mice, that point appears to be before the terminal differentiation of photoreceptors. In humans, photoreceptor maturation occurs in central-to-peripheral order and over a period of 4-5 years, beginning at mid-gestation (62-66). Most photoreceptors are already post-mitotic and undergoing terminal differentiation at birth. Translating the results from *Lztfl1* mutant mice to human gene therapy, our findings suggest that therapeutic genes must be delivered within the first 4-5 years in life to secure the maximal and long-term benefits of gene therapy. This is not simply because of the number of photoreceptors viable at the time of treatment. Instead, our data suggest that there is a qualitative difference between immature and mature photoreceptors (or retinas) with regard to their capabilities to recover, and this capability declines rapidly as photoreceptors undergo maturation. Therefore, our study also highlights the importance of early diagnosis.

Finally, our study raises several important questions. Which diseases are amenable to gene therapy even at late disease stages and which ones need very early intervention to reap long-lasting benefits? If gene therapy does not have a permanent effect, what factors limit the permanent rescue of photoreceptors? And how can we overcome such obstacles? Related to the latter question, it was shown that treatment of the retina with ciliary neurotrophic factor (CNTF) prior to the therapeutic gene delivery in *CNGB3*-associated achromatopsia caused transient dedifferentiation of photoreceptors and significantly improved the efficacy of late stage gene therapy (67). Similar strategies might be developed. Future studies should address these questions.

## MATERIALS AND METHODS

### Animal study approval

This study was carried out in accordance with the recommendations in the Guide for the Care and Use of Laboratory Animals of the National Institutes of Health. All animal studies were approved by the Institutional Animal Care and Use Committee (IACUC) of the University of Iowa.

### Mouse and genotyping

The *Lztfl1*^*gt*^ mouse line was previously described (30). To introduce TMX-inducible FLP recombinase, *Lztfl1*^*gt/gt*^ mice were crossed to B6N.129S6(Cg)-*Gt(ROSA)26Sor*^*tm3(CAG-flpo/ERT2)Alj*^/J mice (The Jackson Laboratory; stock number 019016; (39)). The excision efficiency of the gene-trap was significantly higher in homozygous *FLP*^+^*/FLP*^+^ mice compared with heterozygous *wt/FLP*^+^ mice, and all *Lztfl1*^*gt/gt*^;*FLP*^+^ mice used for rescue experiments were *FLP*^+^*/FLP*^+^. *FLP*− refers to *wt/wt. Ift88*^*fl/fl*^ and *Bbs8*^*gt/gt*^ mice are from our colony and previously described (31, 68). *Rhodopsin-Cre* (also known as *iCre75*) transgenic mice were from Dr. Ching-Kang Chen (Baylor College of Medicine) (69). All animals were maintained under a 12-hr light/12-hr dark cycle with free access to food and water. Mouse tail snips were collected at the time of weaning (P19-P24), and genomic DNAs were extracted by Proteinase K digestion (Sigma-Aldrich; RPROTK-RO) in Tail Lysis Buffer (10 mM Tris pH 8.0, 100 mM NaCl, 10 mM EDTA, 1% SDS, 0.3 mg/ml Proteinase K) followed by ethanol precipitation. Genotyping was conducted by PCR using GoTaq G2 Flexi DNA polymerase (Promega). Primers used for genotyping are listed in **Table S1**. All primers were obtained from Integrated DNA Technologies (IDT). PCR protocols are available upon request.

### Tamoxifen (TMX) administration and genomic DNA PCR for excision efficiency tests

TMX (Sigma-Aldrich; T5648) was dissolved in corn oil (Sigma-Aldrich; C8267) to a concentration of 40 mg/ml. Mice at P21 and older were intraperitoneally injected with TMX at a dose of 120 mg/kg body weight for 5 consecutive days (1 injection/day). For P5/8/12 injections, mice were intraperitoneally injected with 0.5 mg (at P5), 0.8 mg (at P8), and 1 mg (at P12) of TMX (1 injection/day), respectively. To assess excision efficiencies after TMX injection, genomic DNAs were extracted from mouse tail snips as described above and used for PCR analyses. To assess excision efficiencies in photoreceptors, the ONL was isolated by laser-capture microdissection. Briefly, mice were euthanized by CO2 asphyxiation followed by cervical dislocation, and eyes were enucleated and frozen in Neg-50 Frozen Section Medium (Richard-Allan Scientific). Ten μm cryosections were collected on PEN membrane glass slides (Applied Biosystems) using a CryoStar NX70 Cryostat (Thermo Scientific). Sections were fixed with 70% ethanol for 30 sec followed by dehydration in 95% and 100% ethanol (30 sec each). After drying, the ONL was micro-dissected from retinal sections using LMD7000 Laser Capture Microdissection System (Leica). Genomic DNAs were extracted as described above using Proteinase K and Tail Lysis Buffer. PCR primers used for excision efficiency tests are listed in **Table S1**.

### ERG

Mice were dark-adapted overnight before ERG recording. Under dim red light illumination, mice were anesthetized with intraperitoneal injection of ketamine and xylazine (87.5 mg/kg and 12.5 mg/kg, respectively), and pupils were dilated with 1% tropicamide ophthalmic solution (Akorn) for 2-3 minutes. Mice were placed on a Celeris D430 rodent ERG testing system (Diagnosys) with its heater on to maintain animal’s body temperature. After applying a drop of GenTeal Tears Lubricant Eye Gel (Alcon), light-guide electrodes were positioned on both eyes. Dark-adapted Standard Combined Response (SCR) was measured with 15 flashes of 3.0 cd×sec/m^2^ stimulating light (color temperature: 6500K). For photopic ERG, mice were light-adapted for 10 minutes (background light; 9.0 cd×sec/m^2^) and responses were measured with 15 flashes of 3.0 cd×sec/m^2^ white light (6500K). After recording, animals were allowed to recover on a heating pad. Statistical significance was determined by one-way ANOVA with Tukey’s multiple comparison test using GraphPad Prism (p<0.01).

### RNA extraction and qRT-PCR

Total RNAs were extracted from eyes using Polytron PT 1200E homogenizer (Kinematica) and TRI Reagent (Sigma) following the manufacturer’s instructions. One μg of RNA was used for cDNA synthesis with random hexamer primers (Invitrogen) and SuperScript III reverse transcriptase (Invitrogen). Quantitative PCR was conducted using SsoAdvanced Universal SYBR Green Supermix (Bio-Rad) and CFX96 Real-Time PCR Detection System (Bio-Rad). *Rpl19* mRNA levels were used for normalization and the ΔΔCt method (70) was used to measure relative gene expression levels. Production of a single PCR product from each primer pair was validated by melt-curve analyses, and the amplification of correct fragments was confirmed by Sanger sequencing. Primer sequences are described in **Table S1**.

### Protein extract preparation, SDS-PAGE, and Immunoblotting

Mice were euthanized by CO2 asphyxiation followed by cervical dislocation. Eyes were enucleated and either used directly for preparing whole eye protein extracts or neural retinas were dissected for retinal protein extracts. Collected tissues were homogenized with Polytron PT 1200E in Lysis Buffer (50 mM HEPES pH 7.0, 150 mM NaCl, 2 mM EGTA, 2 mM MgCl2, 1% Triton X-100) supplemented with Protease Inhibitor Cocktail (Bimake). Homogenates were centrifuged at 20,000 x*g* for 15 min at 4 °C to remove insoluble materials. Protein concentrations were measured using a DC Protein Assay kit (Bio-Rad) with BSA protein standards and AccuSkan GO spectrophotometer (Fisher Scientific). Mixed with NuPAGE LDS Sample Buffer (Invitrogen) and Reducing Agent (Invitrogen), 50 μg (whole eye extract) or 30 μg (retinal extract) of proteins were loaded per lane on NuPAGE 4-12% (wt/vol) Bis-Tris gels (Invitrogen). Proteins were transferred onto a nitrocellulose membrane (Bio-Rad) and probed with primary antibodies following standard protocols. Primary antibodies used and their specificity/validation data are summarized in **Table S2** and **Figure S9**. For detection, horse radish peroxidase (HRP)-linked anti-mouse IgG or anti-rabbit IgG secondary antibodies (Cell Signaling Technology) were used with SuperSignal West Dura Extended Duration Substrate (Thermo Scientific). Images were taken with a ChemiDoc system (Bio-Rad) and quantified with the Image Lab software (Bio-Rad).

### Immunohistochemistry and TEM

Tissue preparation, fixation, labeling, and imaging procedures for immunohistochemistry and TEM were described previously (40). For immunohistochemistry, Neg-50 (Richard-Allan Scientific) embedded frozen eye cups were mounted on a cryostat chuck in an orientation that sections became perpendicular to the retinal plane at the central retina. Serial sections were collected from the middle 1/3 of eye cups and used for immunohistochemistry. To assess the photoreceptor cell loss, 3 serial sections that contained the most central part of the retina were selected (within 250 μm from the center), and the number of rows of photoreceptor cell nuclei were counted near the center of the retina (2 locations/section). For fluorescence intensity quantification, 20-25 μm width areas were randomly selected and integrated fluorescence intensities were measured within the IS (from the distal end of the IS to synaptic terminals) and the entire photoreceptor cells using ImageJ. The ratio of fluorescence intensity in the IS over the entire cell was presented. One-way ANOVA followed by Tukey’s multiple comparison test was used for statistical analyses using GraphPad Prism software. *P* values smaller than 0.01 were regarded as statistically significant.

## ACKNOWLEDGEMENTS

We thank Dr. Ching-Kang Chen (Baylor College of Medicine) for generously providing *iCre75* transgenic mice. We also thank Dr. Chantal Allamargot for excellent assistance in electron microscopy, Jacob Hunter for assistance in our mouse colony maintenance and genotyping, and Dr. Edwin Stone for critical reading of the manuscript. This work was supported by National Institutes of Health grants R01-EY022616 and R21-EY027431 (to S.S.) and National Institutes of Health Center Support grant P30-EY025580 to the University of Iowa.

## Conflict of Interest Statement

None declared.

## Abbreviations

AAV: adeno-associated virus
BBS: Bardet-Biedl syndrome,
CMV: cytomegalovirus
DAPI: 4′,6-Diamidine-2′-phenylindole dihydrochloride
ERG: electroretinography
FLP: flippase
FRT: flippase recognition target
IFT: intraflagellar transport
IRD: inherited retinal degeneration
IS: inner segment
LCA: Leber congenital amaurosis
ONL: outer nuclear layer
OS: outer segment
P: post-natal day
PTI: post-tamoxifen injection
qRT-PCR: quantitative reverse transcription polymerase chain reaction
RP: retinitis pigmentosa
SCR: standard combined response
TEM: transmission electron microscopy
TMX: tamoxifen

## Notes

### Competing Interest Statement

The authors have declared no competing interest.

