## supplemental for "Limited time window for retinal gene therapy in a preclinical model of ciliopathy"

**This PDF file includes:**

Supplementary text
Figures S1 to S9
Tables S1 to S2
SI References

### Supplementary Information Text

### Extended Materials and Methods

#### *Sucrose gradient ultracentrifugation*

Mouse eyes were collected and homogenized with Polytron PT 1200E in ice-cold PBS Lysis Buffer (PBS with 0.7% Triton X-100). After centrifugation at 20,000 x*g* for 15 min at 4 °C, supernatants were concentrated with Microcon Centrifugal Filter devices (20,000 MWCO; Millipore), loaded on a 4-ml 10-40% sucrose gradient in PBS with 0.04% Triton X-100, and spun at 166,400 x$g_{avg}$ for 15 hours (Sorvall WX100 Ultra Centrifuge; Thermo Scientific). A total of 19 fractions (~210 µl each) were collected from the bottom of each centrifuge tube using a 26-G needle. Fifteen microliters of each fraction were used for SDS-PAGE and immunoblotting following standard protocols.

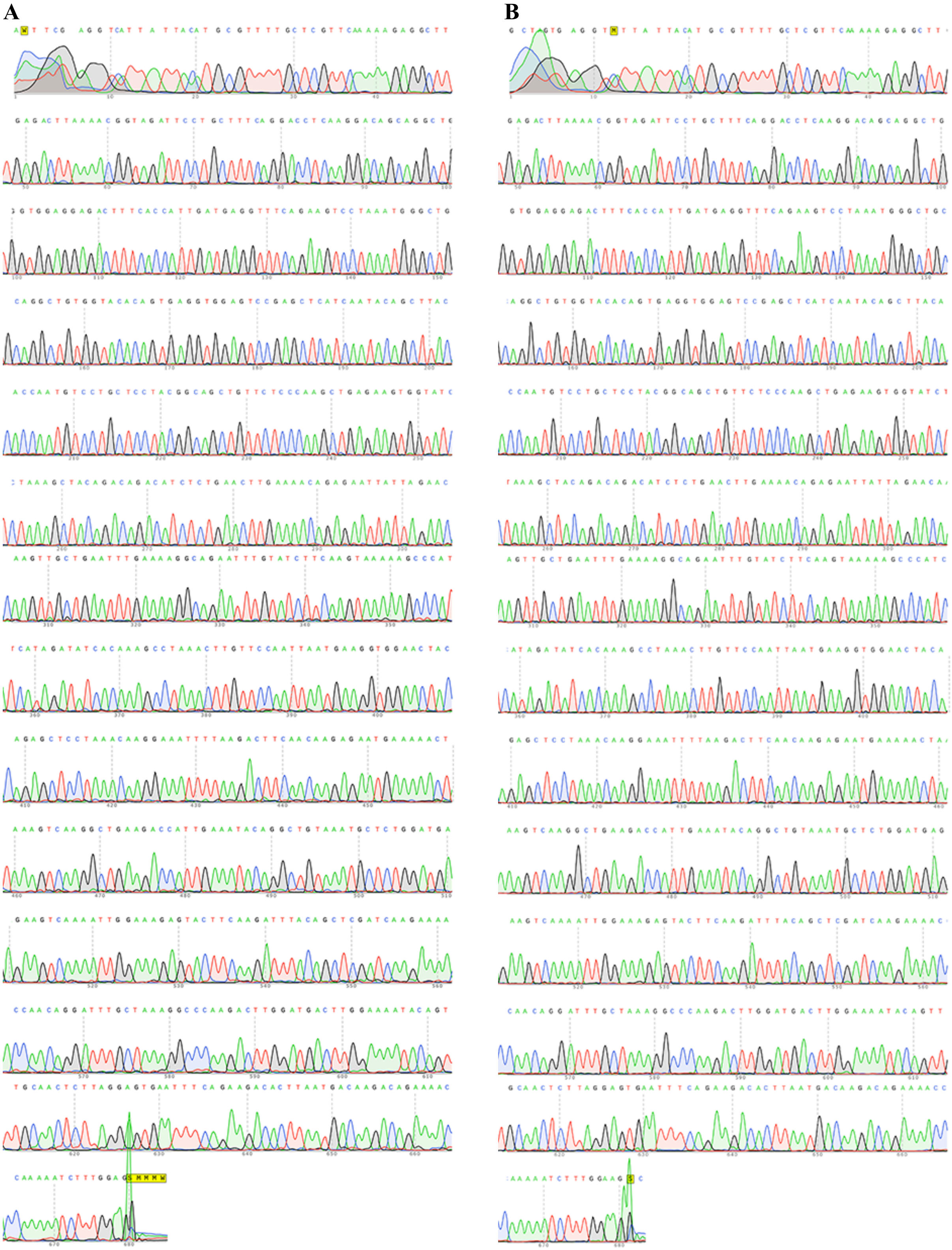

**Fig. S1. Sequencing results of *Lztfl1* cDNAs (between exons 2 and 8)**

A) Sequence of *Lztfl1* cDNA from normal (*Lztfl1*$^{+/gt}$) mouse eyes.

B) Sequence of *Lztfl1* cDNA from rescued *Lztfl1*$^{gt/gt}$ mouse eyes.

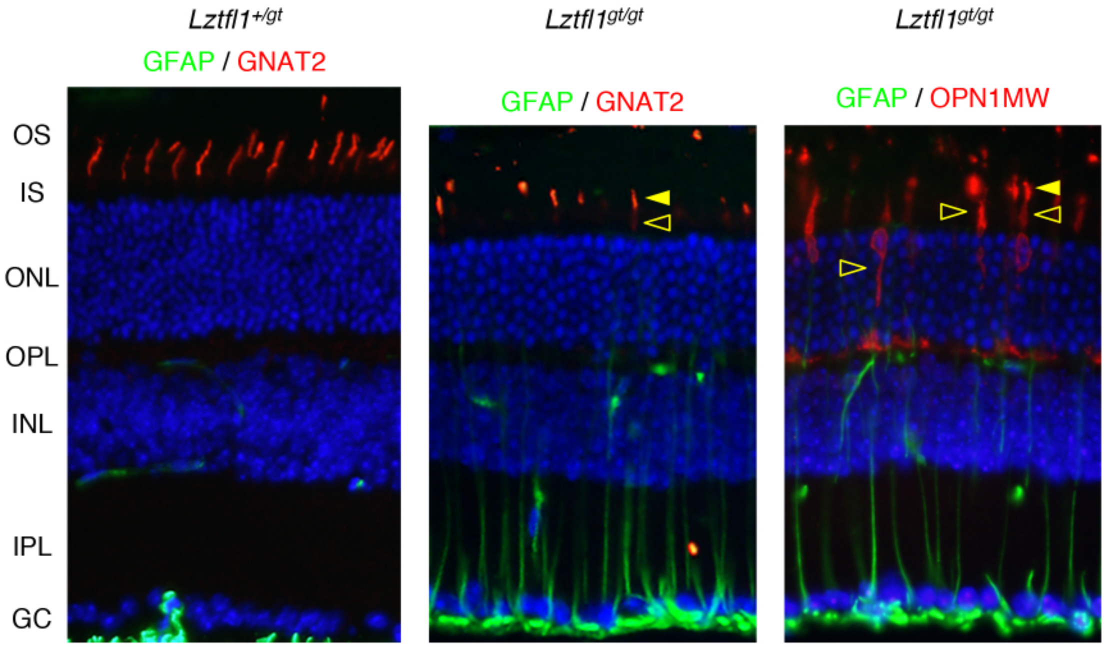


**Fig. S2. Shortening of the cone OS in *Lztfl1* mutant retinas.**

Retinal sections from *Lztfl1*$^{+/gt}$ and *Lztfl1*$^{gt/gt}$ mice at P45 were decorated with GFAP (green), GNAT2 (red), and OPN1MW (red) antibodies. Cone OSs in *Lztfl1* mutant retinas were marked by yellow arrowheads. Mislocalization of OPN1MW to the IS was marked by open arrowheads.

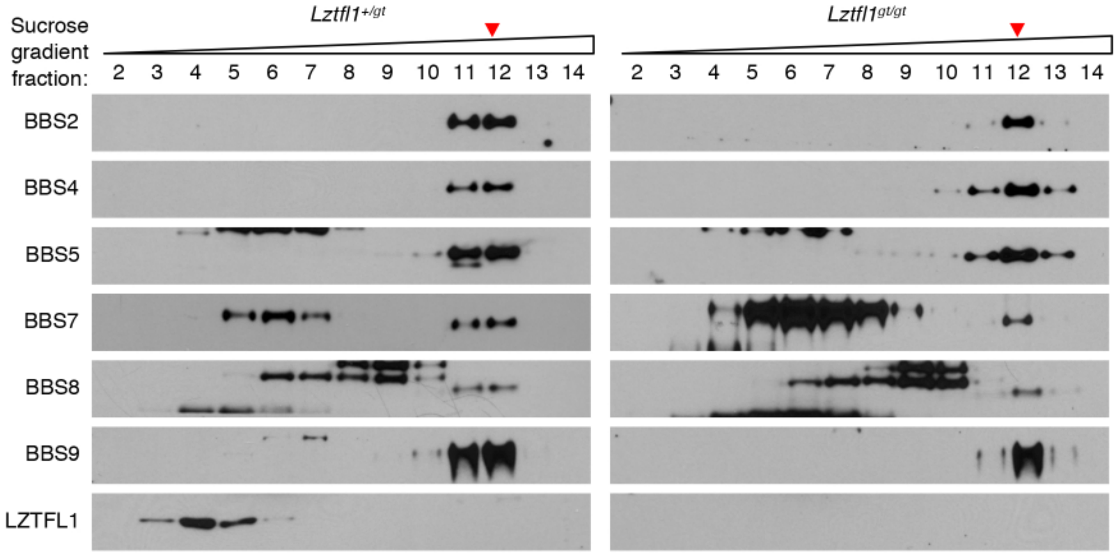


**Fig. S3. BBSome assembly is not altered in *Lztfl1*$^{gt/gt}$ mutant eyes.**

Protein extracts from normal (*Lztfl1*$^{+/gt}$) and *Lztfl1*$^{gt/gt}$ mutant eyes were loaded on 10-40% sucrose gradients and proteins were separated by sucrose gradient ultracentrifugation. Sedimentation of individual BBSome components was examined by SDS-PAGE and immunoblotting. Red arrowheads mark the peak at which the BBSome components were co-fractionated.

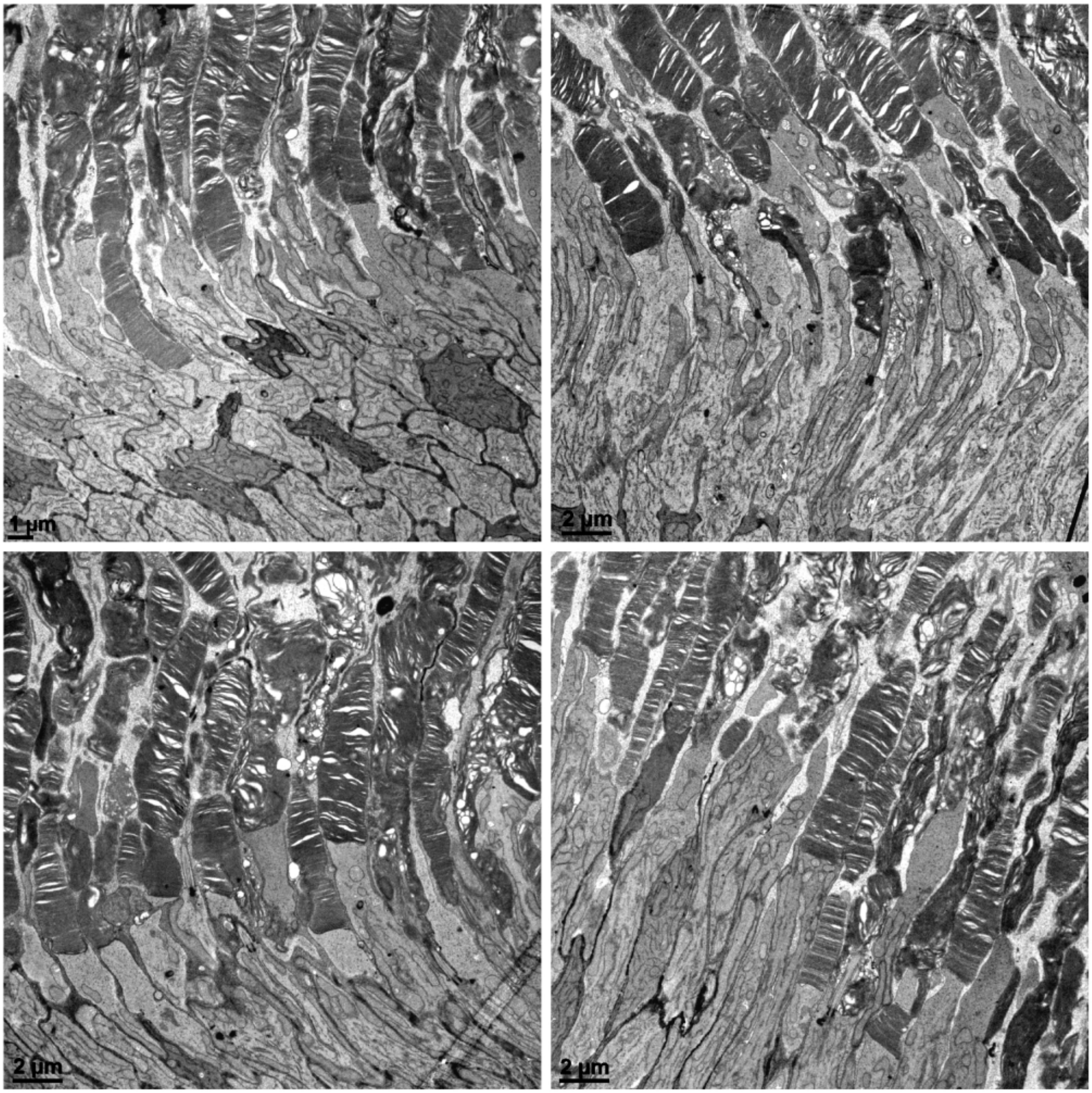


**Fig. S4. TEM images of P5-rescued *Lztfl1*$^{gt/gt}$*;FLP*$^{+}$ mice at 3 months PTI.**

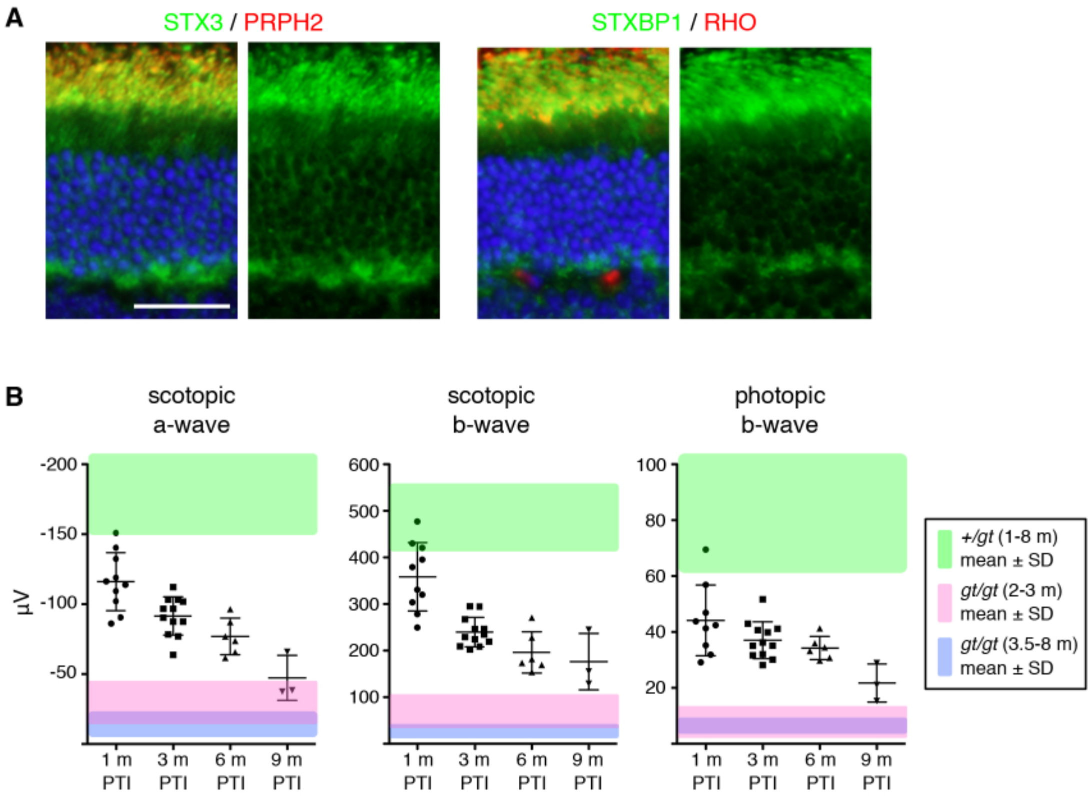


**Fig. S5. Protein mislocalization and ERG in P5-rescued *Lztfl1$^{gt/gt}$;FLP$^{+}$* mice at 9 months PTI.**

A) Retinal sections from P5-rescued *Lztfl1$^{gt/gt}$;FLP$^{+}$* mice were collected at 9 months PTI and stained with anti-STX3 (green) and anti-STXBP1 (green) antibodies. Anti-PRPH2 (red) and anti-RHO (red) antibodies were used as a marker of the OS. Merged images are shown on the left. Scale bar denotes 25 µm.

B) ERG responses of P5-rescued *Lztfl1$^{gt/gt}$;FLP$^{+}$* mice at 9 months PTI. Data were added to the graphs presented in Figure 3 for direct comparisons with prior time points (1 m, 3 m, and 6 m PTI). Statistical analyses were not conducted because of the small number of animals examined.

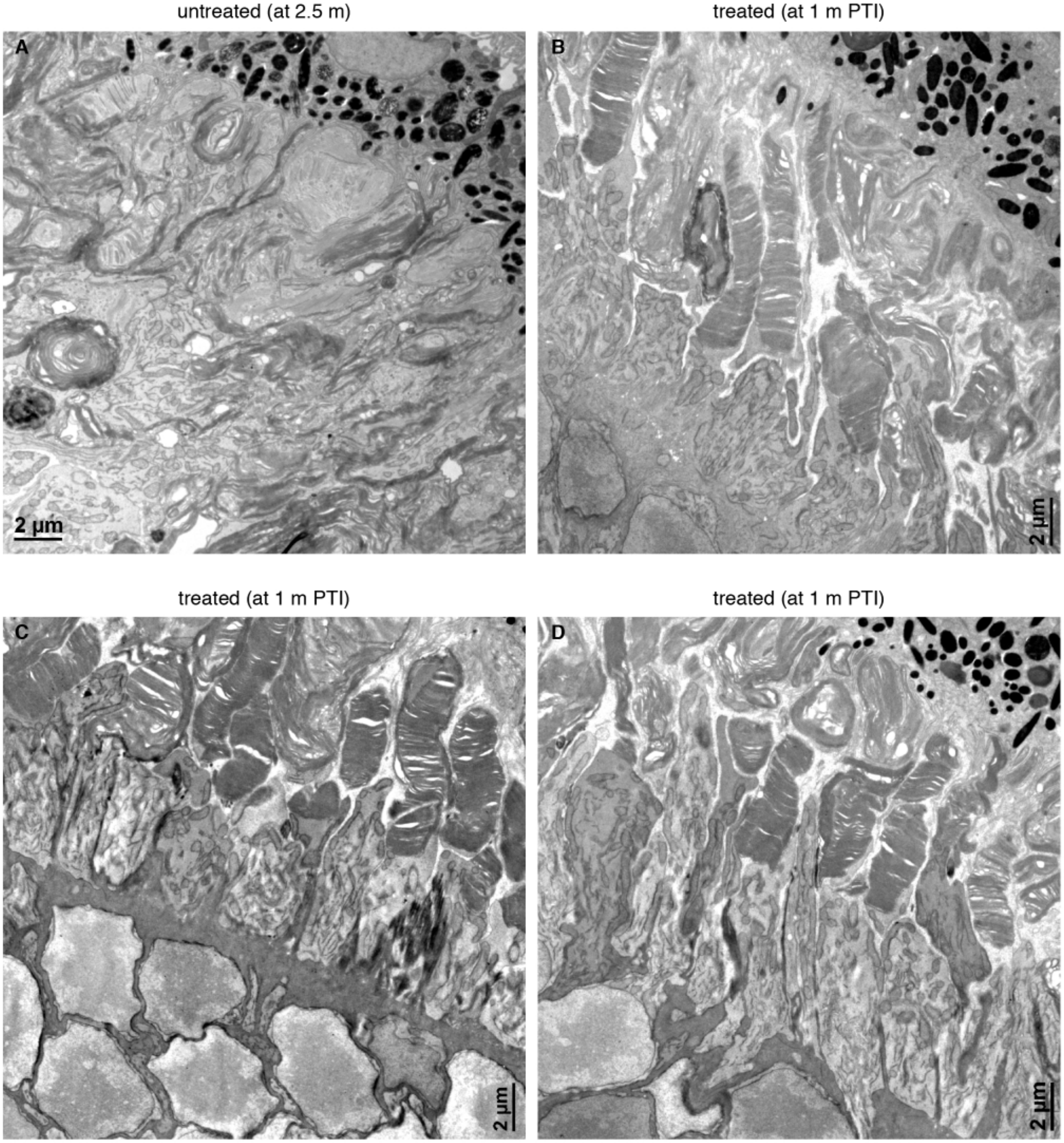


**Fig. S6. TEM images of P45-rescued *Lztfl1*$^{gt/gt}$*;FLP*$^{+}$ mice at 1 month PTI.**

Transmission electron micrographs from untreated (A; 2.5-months old) and P45-treated (B-D) *Lztfl1*$^{gt/gt}$*;FLP*$^{+}$ mice at 1 month PTI.

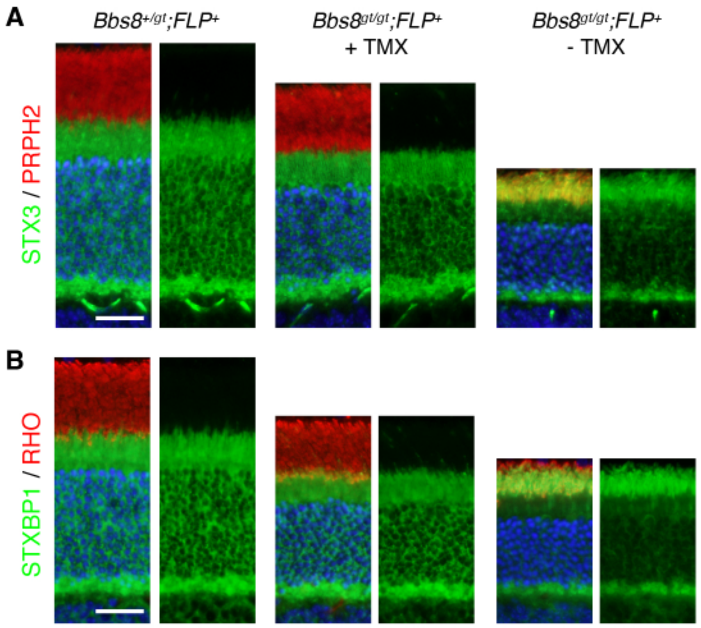


**Fig. S7. Restoration of *Bbs8* expression at P25-29 completely rescues the protein mislocalization phenotype in *Bbs8*$^{gt/gt}$ mice.**

Retinal sections from P25-rescued *Bbs8*$^{gt/gt}$*;FLP*$^{+}$ mice (middle) were collected at 1.5 months PTI and stained with STX3 (A; green) and STXBP1 (B; green) antibodies. Retinas from age-matched (2.5 months old) normal (*Bbs8*$^{+/gt}$*;FLP*$^{+}$; left) and untreated *Bbs8*$^{gt/gt}$*;FLP*$^{+}$ (right) mice were included as controls. PRPH2 (red) and RHO (red) antibodies were used as a marker of the OS. Scale bar denotes 25 μm.

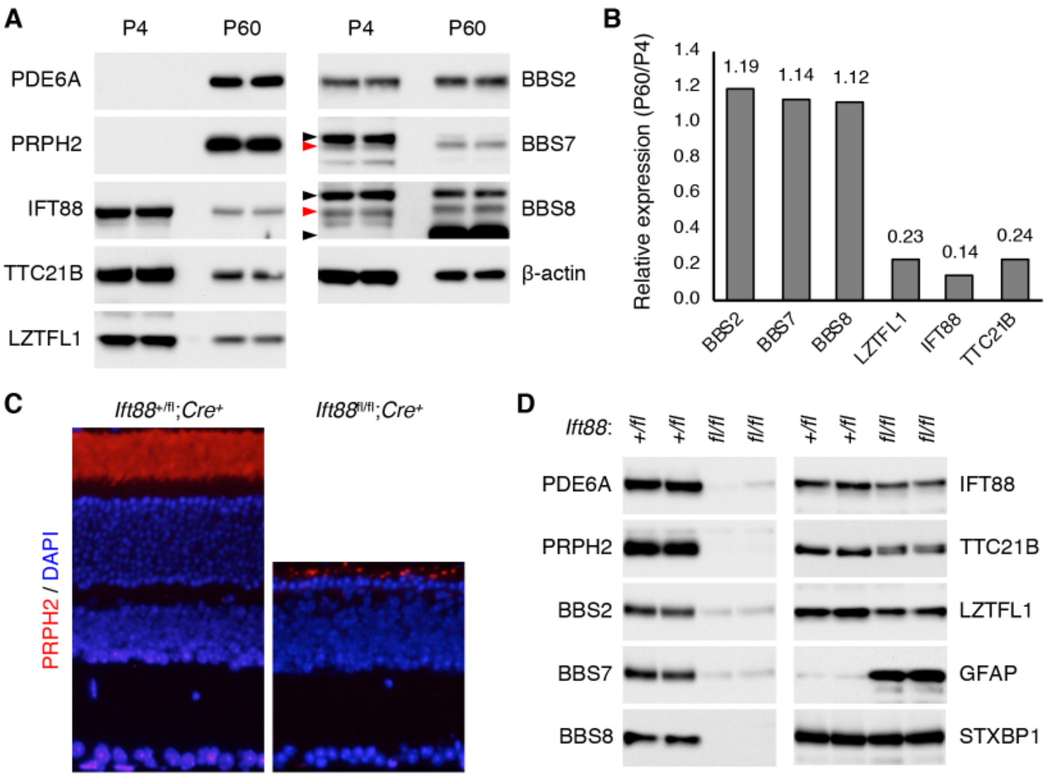


**Fig. S8**. **Expression of LZTFL1 is more similar to IFT proteins than BBSome components.**

A) Retinal protein extracts from 4-day and 60-day old wild-type mice were analyzed by immunoblotting. Red arrowheads mark BBS7 and BBS8 proteins, while black arrowheads indicate cross-reacting proteins.

B) Relative expression levels of BBS2, BBS7, BBS8, LZTFL1, IFT88, and TTC21B at P60 compared with at P4.

C) Retinal sections of 2-month old *Ift88*$^{+/fl}$*;rhodopsin-Cre*$^{+}$ and *Ift88*$^{fl/fl}$*;rhodopsin-Cre*$^{+}$ mice. More than 95% of photoreceptors are lost by this age.

D) Retinal protein extracts from 2-month old *Ift88*$^{+/fl}$*;rhodopsin-Cre*$^{+}$ and *Ift88*$^{fl/fl}$*;rhodopsin-Cre*$^{+}$ mice were analyzed by immunoblotting.

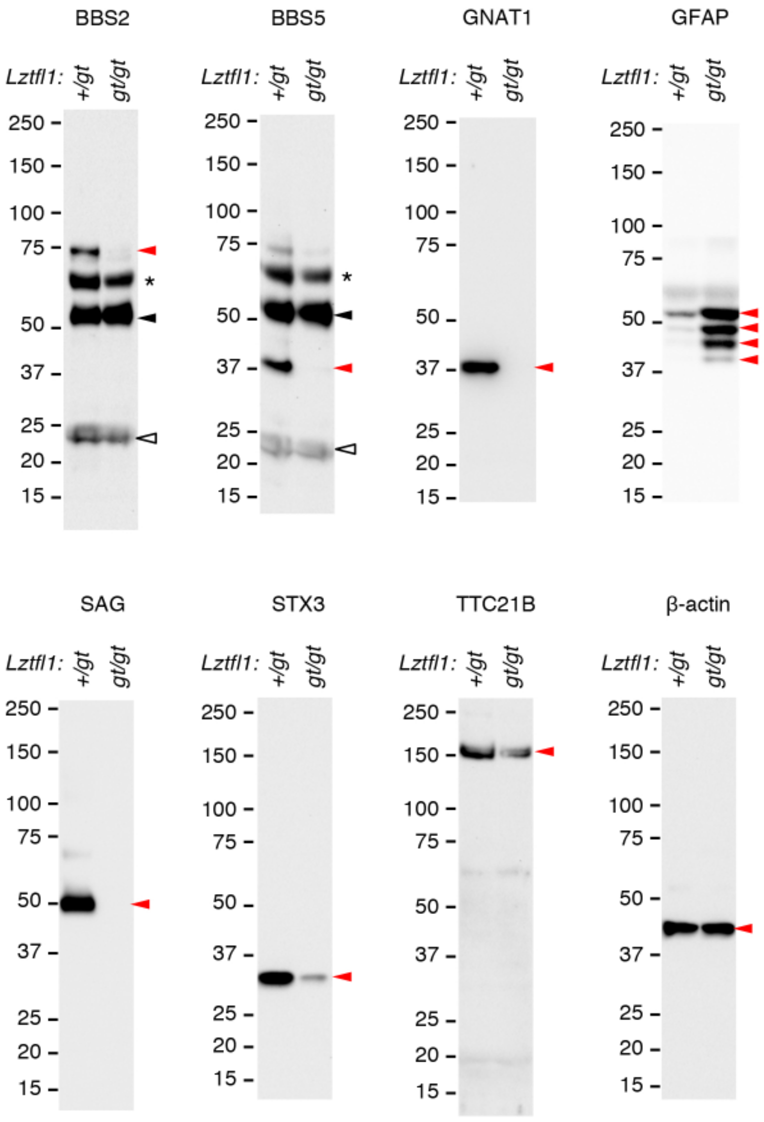


**Fig S9. Specificity of the antibodies used in this study.**

Whole eye protein extracts from 6-month old normal (*Lztfl1*$^{+/gt}$) and "photoreceptor-less" *Lztfl1*$^{gt/gt}$ mice were separated on an SDS-PAGE gel, and indicated antibodies were used for immunoblotting. Target protein bands were marked by red arrowheads. Black and open arrowheads indicate IgG heavy and light chains, respectively. Asterisks denote unknown, cross-reacting proteins.

**Table S1. List of PCR primers used**

| Purpose | Primer Name | Sequence | Product size |
|---|---|---|---|
| *Lztfl1$^{gt}$* genotyping | F-Lztfl1-comm | TAACATGCCACTTGGACATCATGG | wt: 409 bp<br>gt: 543 bp |
| | R-Lztfl1-wt | ATTCCATGAAAGCTGGTGTTGTGA | |
| | R-Lztfl1-gt | CCACAACGGGTTCTTCTGTTAGTC | |
| *FlpER* genotyping | F-FLP-comm | AAAGTCGCTCTGAGTTGTTAT | wt: 603 bp<br>FLP: 309 bp |
| | R-FLP-wt | GGAGCGGGAGAAATGGATATG | |
| | R-FLP-tg | TTATGTAACGCGGAACTCCA | |
| *Ift88* genotyping | F-Ift88-flox | GACCACCTTTTTAGCCTCCTG | wt: 209 bp<br>fl: 254 bp |
| | R-Ift88-flox | TTCTGGCTCTGAACACAATCC | |
| *iCre75* genotyping | F-iCre75 | TCAGTGCCTGGAGTTGCGCTGTGG | wt: none<br>Tg: 650 bp |
| | R-iCre75 | CTTAAAGGCCAGGGCCTGCTTGGC | |
| *Lztfl1$^{gt}$* excision | F-Lz-exc-comm | GCGCATAACGATACCACGATA | unexcised: 599 bp<br>excised: 292 bp |
| | R-Lz-unexcised | AGAGACAGGTGAGAGGAGATG | |
| | R-Lz-excised | GGTACAGCAGTGATTTCCCTATT | |
| *Lztfl1* exon 2 qRT-PCR | F-Lz-exon2 | GGCCTAAATGAGCACCATCA | 103 bp |
| | R-Lz-exon2 | GGTCCTGAAAGCAGGAATCTAC | |
| *Lztfl1* exon 10 qRT-PCR | F-Lz-exon10 | CCCAAATACGCCTCTGTCAT | 95 bp |
| | R-Lz-exon10 | GGCTCTTCTTCCCACTCTAAAC | |
| *Lztfl1* cDNA (exons 2-8) | F-Lz-RT2 | GCAGAGTTGGGCCTAAATGA | 722 bp |
| | R-Lz-RT8 | GCTGCTAGGTTCTCTTCCAAA | |
| *Rpl19* qRT-PCR | F-Rpl19-qRT | GCAAGCCTGTGACTGTCCATT | 106 bp |
| | R-Rpl19-qRT | GCATTGGCAGTACCCTTCCTC | |

**Table S2. List of antibodies used**

| Antibody | Source / Reference | Cat# | Verification |
|---|---|---|---|
| Mouse anti-β-actin monoclonal (clone: AC-15) | Sigma-Aldrich | A1978 | (1), this study |
| Mouse anti-BBS2 monoclonal (clone: A-12) | Santa Cruz | sc-365355 | this study |
| Rabbit anti-BBS4 polyclonal | Maxence Nachury (2) | N/A | (2, 3) |
| Mouse anti-BBS5 polyclonal (clone: B-11) | Santa Cruz | sc-515331 | this study |
| Rabbit anti-BBS7 polyclonal | ProteinTech Group | 18961-1-AP | (4) |
| Rabbit anti-BBS8 polyclonal | Sigma-Aldrich | HPA003310 | (4) |
| Rabbit anti-BBS9 polyclonal | Sigma-Aldrich | HPA021289 | (3) |
| Mouse anti-GFAP monoclonal (clone: GA5) | EMD Millipore | MAB3402 | (5), this study |
| Rabbit anti-GNAT1 polyclonal | ProteinTech Group | 55167-1-1AP | this study |
| Rabbit anti-GNAT2 polyclonal | abcam | ab97501 | (6) |
| Rabbit anti-IFT88 polyclonal | ProteinTech Group | 13967-1-1AP | (7, 8) |
| Rabbit anti-LZTFL1 polyclonal | Seongjin Seo (3) | N/A | (3, 6) |
| Rabbit anti-PDE6A polyclonal | ProteinTech Group | 21200-1-1AP | (6) |
| Rabbit anti-PRPH2 polyclonal | ProteinTech Group | 18109-1-AP | (6) |
| Mouse anti-RHO monoclonal (clone: 1D4) | EMD Millipore | MAB5356 | (9) |
| Mouse anti-SAG monoclonal (clone: E-3) | Santa Cruz | sc-166383 | this study |
| Mouse anti-STX3 monoclonal (clone: 1-146) | EMD Millipore | MAB2258 | this study |
| Rabbit anti-STXBP1 polyclonal | ProteinTech Group | 11459-1-AP | (6) |
| Rabbit anti-TTC21B polyclonal | Sigma-Aldrich | HPA035495 | this study |
